# Cellularization in chytrid fungi uses distinct mechanisms from conventional cytokinesis and cellularization in animals and yeast

**DOI:** 10.1101/2025.01.30.635136

**Authors:** Edgar M. Medina, Mary Williard Elting, Lillian Fritz-Laylin

## Abstract

Chytrid fungi provide a model for studying foam-like cellularization, where nuclei that are dispersed throughout the cytoplasm are synchronously compartmentalized into daughter cells. This organization poses geometric challenges not faced by cells undergoing conventional cytokinesis or *Drosophila* monolayer cellularization, where nuclei are organized in linear or planar arrangements with ready-access to the plasma membrane. We use the chytrid *Spizellomyces punctatus* to show that chytrid cellularization begins with migration of nuclei and their attached centrosomes to the plasma membrane, where centrosome-associated vesicles mark sites of membrane invagination. These vesicles then extend inwards, resulting in tubular furrows that branch and merge to create a honeycomb of polyhedral membrane compartments–a cellularization foam–each with a nucleus and cilium. Using inhibitors and laser ablation, we show that tensile forces produced by actomyosin networks drive aphrogenesis (foam-generation), while microtubules are important for foam patterning and ciliogenesis but are not essential for cellularization. Finally, we suggest that chytrids may have incorporated ancestral mechanisms associated with ciliogenesis to coordinate the association of internal nuclei with membrane furrows to solve the unique geometric challenges associated with aphrogenic cellularization.

## Introduction

Animal and fungal cells typically divide by forming a trench-like membrane furrow that deepens to separate daughter nuclei into two distinct compartments. ^1,2^ This “conventional” cytokinesis occurs after a single round of nuclear division with the two daughter nuclei arranged in-line. When multiple rounds of nuclear division precede cytokinesis, however, the resulting multinuclear state must be resolved by more complex cellularization processes. During *Drosophila* development, for example, hundreds of nuclei arrange themselves in a monolayer just under the plasma membrane, making the outcome of this “in-plane” cellularization a sheet of cells at the surface of the embryo. ^3^ In contrast, chytrid fungi take the geometry of cellularization to the third dimension. The lifecycle of chytrid fungi, which include important aquatic pathogens and key players in global carbon cycling, involves a cell with dozens of nuclei cellularizing to produce uniform, uninucleated flagellate daughter cells (**Figure 1**).^4–9^ Unlike *Drosophila* and other species that undergo in-plane cellularization, the nuclei of cellularizing chytrids are distributed throughout the cytoplasm. The mechanisms driving this geometrically complex form of cellularization remain unclear.

**Figure 1.**
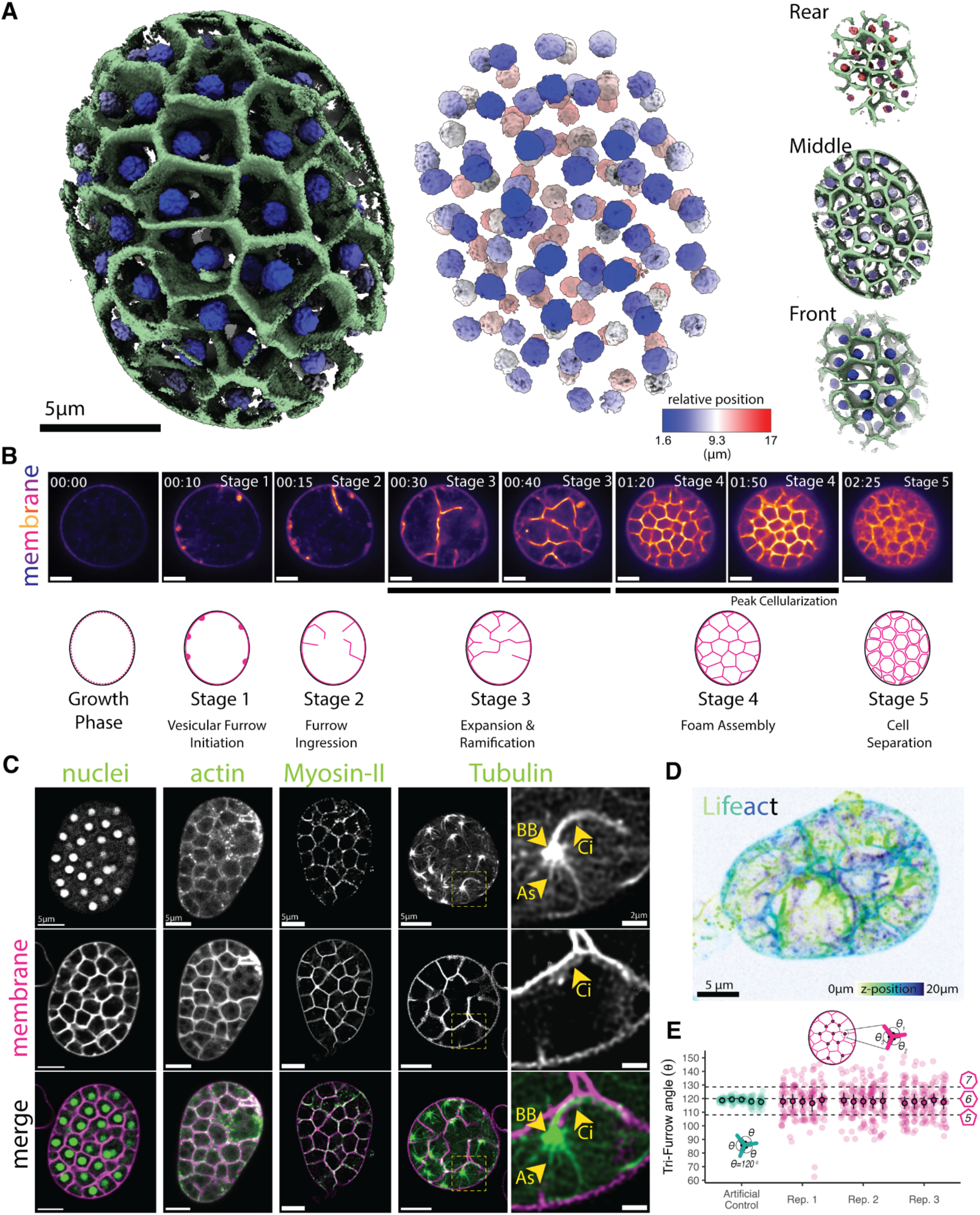
The chytrid *Spizellomyces* cellularizes by building a 3D tessellation of membrane polyhedra of homogeneous size, hexagonal symmetry, and demarcated by actin and myosin-II activity. (A) ChimeraX 3D reconstruction from a point scanner confocal live-cell image of a chytrid sporangium at peak cellularization (Stage 4). Membrane polyhedra are uninucleated and homogeneously distributed across the volume of the sporangium (front, middle, rear). FM4-64 dye was used for membrane (green) and nuclear-localized mClover3 for highlighting nuclei (blue to red). The nuclear position is shown as relative depth from the first stack image. **(B)** Images and idealized schematic of the five main stages of development of the membrane structures of chytrid cellularization. Note that during Stage 5—cell separation—cells remain packed and thus do not acquire a spherical shape until some cells escape the mother. **(C)** Actin and myosin-II colocalize with the membrane of the cellularization polyhedra. Live-cell confocal imaging of strains expressing the (nuclei) NLS-mClover3 to track nuclei (same image used in 3D reconstruction in A), the F-actin probe Lifeact-mClover3 (actin), the myosin-II probe MRLC-mClover3 (MRLC), and mClover3-α-Tubulin (Tubulin) to track microtubules. The magnified region of microtubules highlights the ciliary axoneme (Ci) that starts elongating after entry into cellularization, region containing the basal body (BB), and the astral microtubules and bundles (As). Membrane dyed with FM4-64. Scale 5 µm. **(D)** During mid cellularization, actin (and its associated membrane) form an heterogenous network of surfaces of complex 3D organization and geometry. Image shown is a serial Z-section at 0.39 µm intervals of a sporangium expressing the F-actin probe lifeact-mClover3 during early cellularization, color-coded by depth. **(E)**The cellularization polyhedra have internal vertices with angles with a mean that is in agreement with hexagonal symmetry (magenta) Since regular hexagons cannot form a polyhedron or tile a volume, faces with other symmetries such as pentagons and heptagons may be used to form the 3D tessellation and fill the volume of the mother cell. The angles were measured for at least twenty vertices within the central slice of a sporangium and for five sporangia in three biological replicates. These angles were compared to a set of five artificial vertices (teal) with perfect hexagonal symmetry (120 degrees) and similar membrane thickness and proportions to assess technical measurement error. See also Figure S1 and Video S1-S3.

Despite their different geometries, “conventional” cytokinesis and “in-plane” cellularization involve the same fundamental steps. First, nuclei find their positions. Second, the division site is selected. Third, the cell builds a cytokinetic structure at the division site. Fourth, the cytokinetic structure separates the mother cell into daughter cells. Animal and yeast cells use the same general strategies to complete these four steps. Both lineages rely on the pushing and pulling forces of microtubules to position their nuclei. ^10–12^ Both use plasma membrane-anchored cues to define the position of the cytokinetic furrow. ^1,2^ Finally, both drive inward ingression of a trench-like furrow using a contractile ring made of actin and myosin-II. ^1,2^ These same strategies, however, cannot be extrapolated directly to chytrid cellularization for two reasons. First, interior nuclei are obstructed from access to cues placed on the plasma membrane. Second, it is not obvious how actomyosin rings assembled at the plasma membrane could generate the many planes needed to surround internal nuclei. Although organisms that undergo complex cellularization are common across many eukaryotic lineages,^13–15^ ^16^ it remains unclear whether and how the mechanisms of conventional cytokinesis and in-plane cellularization apply to these types of “volumetric” cellularization.

Understanding the mechanisms driving complex cellularization requires new, genetically tractable model systems. Here, we use our recently-developed genetic tools to dissect the molecular mechanisms driving geometrically complex cellularization in the model chytrid fungus *Spizellomyces punctatus* (*Sp*).^17^ We show that chytrids cellularize by generating a network of membranes with “foam-like” geometry, identify the main stages of cellularization, and use pharmacological and physical perturbations to disrupt them. We find that chytrid cellularization involves a combination of actomyosin machinery similar to that used for conventional cytokinesis along with ciliogenesis-like events not seen in either conventional cytokinesis or *Drosophila* cellularization. These different forms of cellularization illustrate how cells can deploy the same molecular machinery to achieve the same objective through distinct molecular mechanisms.

## Results

### Membranes colocalize with actin and myosin-II to form a three-dimensional cellularization foam

To understand the molecular mechanisms driving chytrid cellularization, we began by determining the spatial distribution of nuclei and membrane during peak cellularization—the stage when the cellularization network fills the cytoplasm of the spherical mother cell, also called a “sporangium”—of the model chytrid *Spizellomyces punctatus* (*Sp*). We grew synchronized cultures of *Sp* adhered to glass imaging dishes and visualized nuclei using the fluorescent protein mClover3 fused to a nuclear-localization signal and staining the membrane with FM4-64. 3D point-scanner confocal imaging showed that during peak cellularization, each homogenously-distributed nucleus is enclosed by a membrane polyhedron (**Figure 1A; Video S1**). Neighboring polyhedra share faces in a tightly packed organization—like soap bubbles inside a container—confirming the complex geometry of chytrid cellularization.

To determine how this geometry forms, we conducted time-lapse microscopy of FM4-64-stained membranes during cellularization and identified five main stages of reorganization (**Figure 1B**). First, small, spherical invaginations form at the plasma membrane **(Stage 1)**. These “furrow initials” extend and elongate inwards toward the center of the cell, forming a tube-like membrane projection we call “tubular furrow” **(Stage 2)**. (*Note we recognize this term is an oxymoron but have chosen to retain the term “furrow” not because of its shape, but because of its historical context in the cytokinetic literature.)* These tubular furrows then invade the cytoplasm, branching and merging as they grow **(Stage 3)**, resulting in a tightly packed polyhedral network that resembles a honeycomb that occupies the entire mother cell **(Stage 4)**. Finally, in a single synchronous event of cell separation, called abscission, all the polyhedra turn into individual daughter cells **(Stage 5)**. These daughters then swim and/or crawl out of one or more discharge pores in the enclosing cell wall, leaving behind the empty shell of their mother. To measure the consistency of cells formed by cellularization, we measured the diameters of the daughter zoospores released and found a coefficient of variation of 17% (**Figure S1**). This variation is similar to the variance in cell size observed in age-matched fission yeast (6%) and budding yeast (17%) cells, ^18^ indicating that chytrid cellularization incorporates tight control of cell size.

We next identified cellular machinery associated with the cellularization polyhedra. Animal and yeast cytokinesis relies on conserved cytoskeletal elements, primarily actin, myosin-II, and microtubules.^1^ To explore the roles of these elements in chytrid cellularization, we visualized the distribution of actin networks by expressing the Lifeact probe fused to mClover3 and staining the membrane with FM4-64. We found that during peak cellularization (Stage 4), actin and membrane co-localize to the cellularization honeycomb (**Figure 1C; Video S2**). We next determined the distribution of a fluorescently-tagged myosin regulatory light chain (MRLC)—a proxy for myosin-II activity. ^19^ We found that, like actin, fluorescent MRLC colocalizes with membrane in the cellularization honeycomb, suggesting a role of contractile actomyosin in chytrid cellularization (**Figure 1C**). Finally, by expressing a fluorescently tagged α-tubulin, we observed three previously described microtubule structures closely associated with each nucleus during peak cellularization: ^20–22^ First, a microtubule structure that emanates from a focus in close apposition to each nucleus. We infer this focus is the centriolar centrosome and basal body. Second, a long, bright and homogenous structure extending from the basal body towards the nearest cellularization plane—the ciliary axoneme. Third, a heterogeneous network of astral microtubules that extends from the basal body and surrounds the nucleus (**Figure 1C**).

To dissect in more detail the geometric patterning formed during peak cellularization, we measured the angles formed at the vertices of the cellularization polyhedra in cell cross sections (**Figure 1E**). Sampling cross sections through a polyhedron gives a distribution of the most common shapes of its faces. For example, cross sections though a triangular prism—which has five faces, three rectangular and two triangular—can create polygons with three, four, or five sides, with the majority of cross sections having four sides. Cross sections through the cellularization honeycomb displayed vertices with three furrows meeting at a mean angle of 117.8 degrees, consistent with polyhedra with pentagonal (108°) and hexagonal (120°) faces.This makes sense because regular hexagons alone cannot form a polyhedron or tile three-dimensional space. This organization is consistent with the geometry of a foam, a concept developed to describe the behavior of soap bubbles. A foam is a collection of surfaces that meet Plateau’s Laws (1873), ^23,24^ which state that: 1) the “bubbles” in the system should have smooth, unbroken surfaces, 2) all points on the same soap film surface have a constant mean curvature, 3) when soap bubble surfaces meet in threes they do so at an angle of 120 degrees—resulting in hexagonal symmetry—and 4) when they meet in fours they do so at an angle of 109.47—tetrahedral symmetry. Because chytrid cellularization involves continuous, smooth cellularization membranes as well as pentagonal- and hexagonal-faced polyhedra, we conclude that its furrow network conforms to the geometric principles of a foam. In this view, the cellularization membranes behave like soap films that minimize surface area while partitioning space to generate a close-packed array of nearly uniform cell volumes.

Finally, to explore how the tubular furrows of early cellularization transform into the cellularization foam, we generated a volumetric image during furrow expansion and ramification (Stage 3) using our actin probe (**Figure 1D; Video S3**). We found that the invaginating tubular furrows form complex, irregular shapes, more akin to stretching melted cheese than the orderly furrows of *Drosophila* cellularization. Together, these results show that, much like animal and yeast cytokinesis, chytrid cellularization uses actin and myosin-II to create cytokinetic structures that are initiated at the plasma membrane. However, unlike animals and yeast, chytrids do not use orderly, trench-like furrows. Instead, chytrids use irregular, tubular furrows to eventually weave a complex space-filling honeycomb made of polyhedral cells of homogenous volume and pentagonal and hexagonal faces—a cellularization foam.

### Nuclear migration to the plasma membrane precedes the onset of cellularization

In conventional animal and yeast cytokinesis, as well as during *Drosophila* cellularization, daughter nuclei remain stationary while cytokinetic furrows extend between or around them. Because nuclei remain highly dynamic during chytrid cellularization, it is unclear if nuclear position plays a role in defining the placement and assembly of furrows. To explore possible coordination between nuclear movement and furrow formation, we imaged synchronized cultures stained for membranes with FM4-64 and highlighting nuclei with a fluorescently tagged histone H2B. Imaging cross-sections through the center of sporangia revealed a possible enrichment of nuclei at the cell surface just before cellularization (**Figure 2A; Video S4**). To quantify this impression, we calculated the ratio of nuclei in two regions of equivalent area: an outer donut at the surface of the sporangium and the remaining inner disk—the cortical nuclear ratio. This ratio was near one before and during cellularization, except for a short window of ten minutes during which nuclei are enriched in the cortical region (**Figure 2A-B**). To test whether peak cortical nuclear ratio coincides with early stages of cellularization, we aligned the ratio of individual sporangia relative to the time of first vesicular furrow formation (Stage 1, **Figure 2B** top) or to the start of furrow elongation (Stage 2, **Figure 2B**, bottom). We found that cortical nuclear enrichment occurs around 10 minutes after the start of Stage 1 (**Figure 2B**, top), and coincides with the start of furrow elongation (**Figure 2B**, bottom), with most sporangia (22 out of 24, 92%) entering furrow elongation (Stage 2) within five minutes of maximum nuclear enrichment (**Figure S1**). Thus, entry into cellularization is marked by the recruitment of nuclei to the plasma membrane.

**Figure 2.**
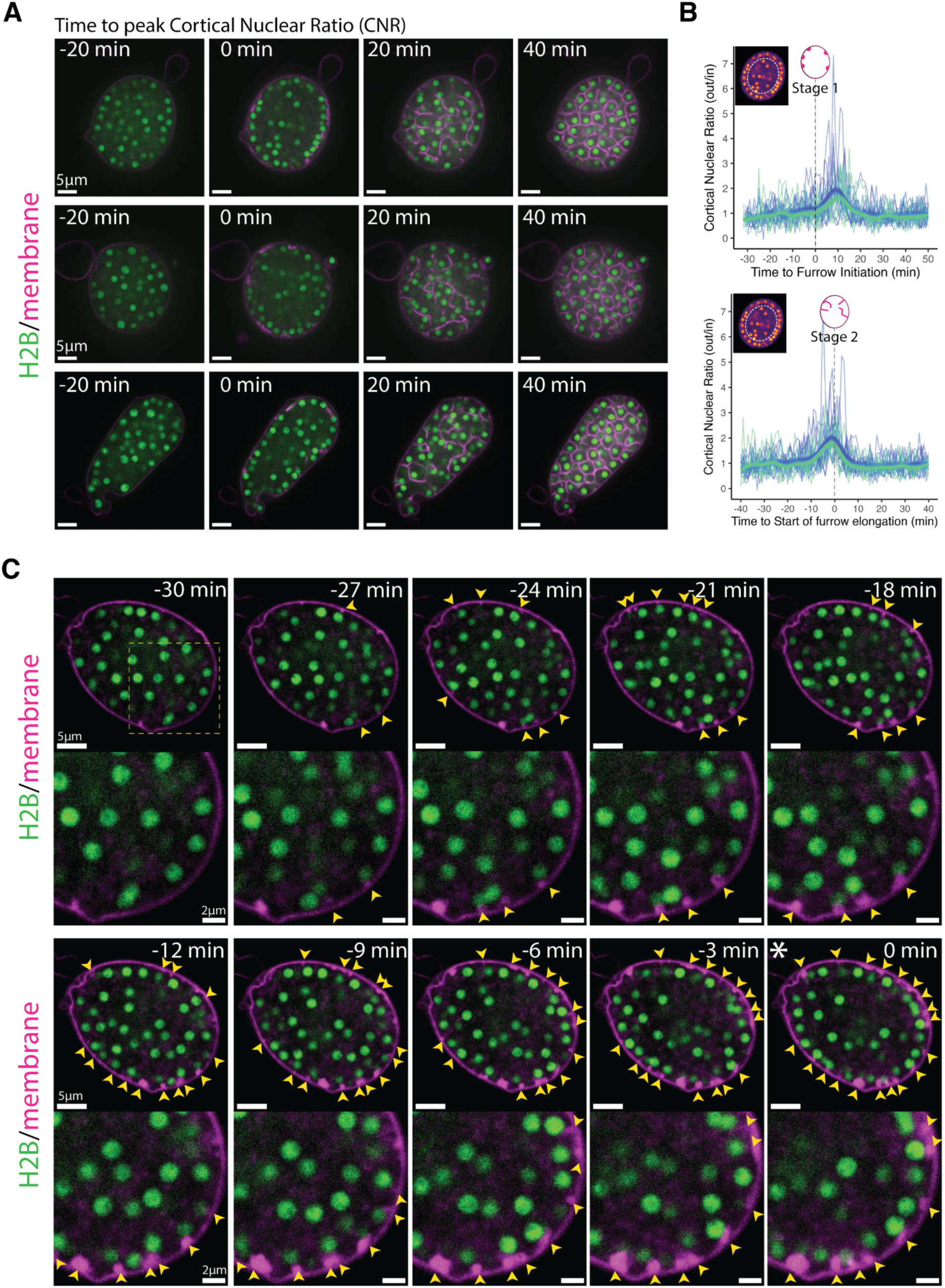
The onset of chytrid cellularization is marked by the recruitment of nuclei to the plasma membrane and the formation of vesicular membrane structures near cortical nuclei. **(A)** Chytrid nuclei (H2B-tdTomato) are homogeneously distributed across the sporangium before (-20 min) and after (+20 min) entry into cellularization except for a narrow window just at entry into cellularization (0 min; asterisk; coincides with the formation of the first membrane vesicles; Stage 1) when nuclei are enriched at the plasma membrane. Scale bars are 5 µm **(B)** Nuclear enrichment at the plasma membrane is limited to a narrow window of time (∼10 min) at the onset of cellularization. After, the nuclei return to be homogeneously distributed. Quantification of nuclear dynamics during time-lapse microscopy 40 min before and after entry into cellularization. The cortical nuclear ratio corresponds to the ratio of the number of nuclei in the outer 50% area (a donut) relative to the inner 50% area (disk)—see inset image. The ratio distribution during time-lapse imaging for each sporangium was centered on zero based on the peak cortical nuclear ratio—marked with an asterisk. Blue and green signify two different biological replicates with thin lines showing individual sporangia and thick lines indicating loess regressions (99% confidence interval; span=0.2). **(C)** Recruitment of nuclei to the plasma membrane during entry into cellularization coincides with the formation of membrane vesicular structures (yellow arrows) in the proximity of each nucleus. Progression of nuclear recruitment and concomitant vesicular furrow initiation shown until peak cortical migration (asterisk; 0 min). Main scale bars 5 µm, inset 2 µm. See also Figure S1, S3, and Video S4.

To further explore the relationship between nuclear position and furrow initiation, we imaged at a higher frame rate and found that vesicular furrow initials form in proximity to the nuclei at the plasma membrane (**Figure 2C**). Imaging of the nuclei approaching entry into cellularization showed a progressive formation of vesicular furrows near cortical nuclei (**Figure 2C**). Because animal cells use astral microtubules to position furrows, we hypothesized that patterning of chytrid cellularization may also involve microtubules. To test this hypothesis we followed cellularization in a strain expressing fluorescent tubulin (mClover3-alpha-Tubulin) in the presence of FM4-64 (**Figure 3A & B; Video S5**). We found that each nucleus in a growing sporangium is associated with two opposing foci of tubulin fluorescence. Because these foci turn into spindle poles and duplicate immediately after nuclear division (**Figure S2A**), we infer that these foci represent centriolar centrosomes. Consistent with a role in furrow positioning for these centrosomes, we observed furrow initials forming in between the centrosomes and the plasma membrane (**Figure 3A & B;Video S5**). To dissect this process in more detail, we built a strain expressing a fluorescently tagged version of SAS-6, a core component of eukaryotic centrioles.^25^ Cells expressing fluorescent SAS-6 show fluorescent foci at spindle poles and the base of the ciliary axoneme consistent with a centriolar distribution. Upon entry into cellularization, SAS-6 puncta approach the plasma membrane, where they become associated with the furrow initials highlighted by FM4-64 (**Figure 3C; Video S6**). The intimate association of centrioles with vesicular furrows agrees with ultrastructural evidence from the cellularization of the zoosporic fungus *Allomyces*, including published electron micrographs showing the presence of basal bodies near cellularization membranes as well as previous observations of nuclear movement coinciding with the start of cellularization (**Figure S3**).^26^ These results suggest that arrival of the centrosomes to the plasma membrane precedes furrow initiation, a finding whose confirmation requires imaging at sub-minute resolution. Unfortunately, such experiments typically result in cells that fail to complete cellularization and die, likely due to phototoxicity. Nevertheless, our results show that during entry into cellularization, nuclei and their associated centrosomes, are transiently recruited to the plasma membrane, where they associate with vesicular furrow initials.

**Figure 3.**
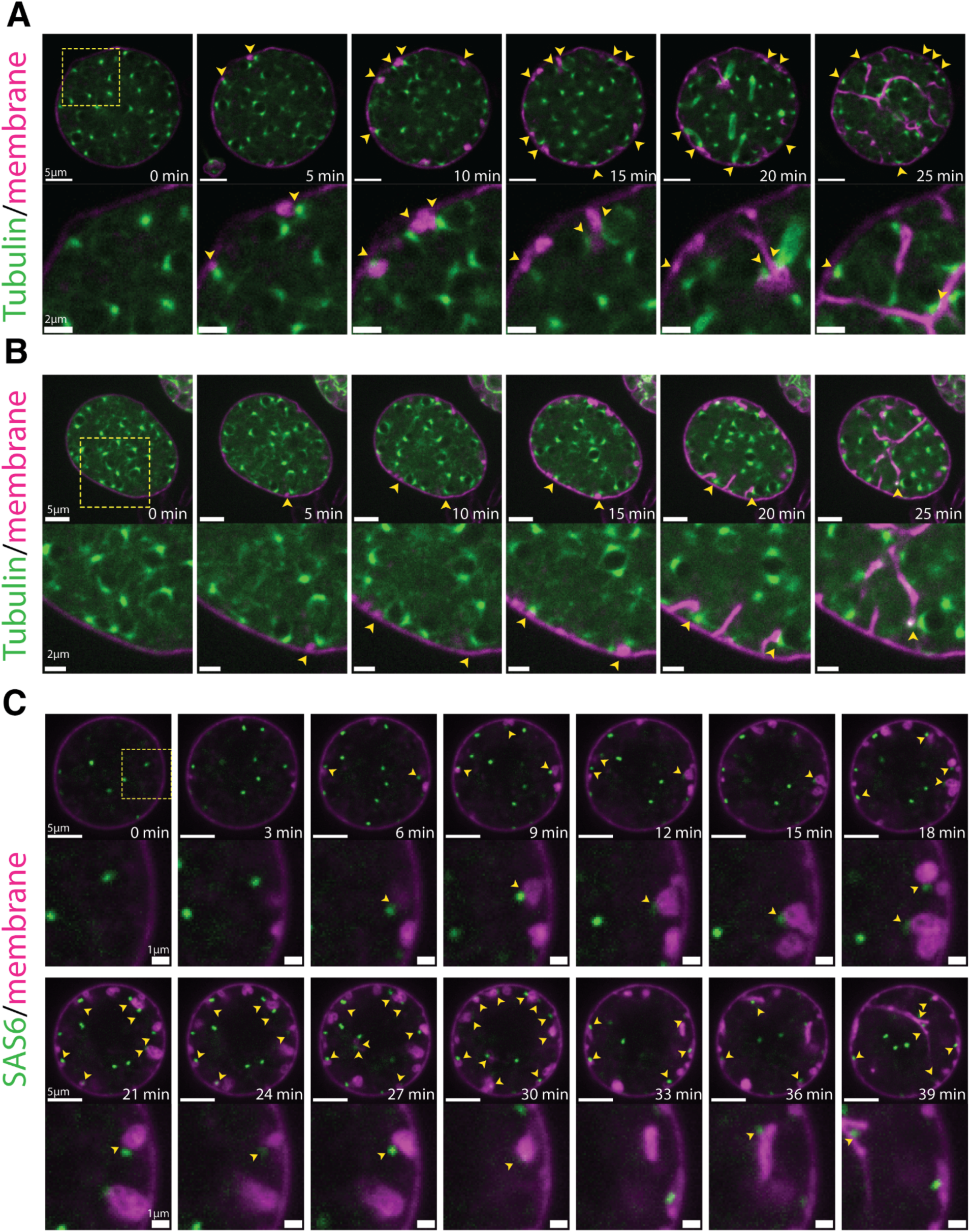
Cortical nuclei attach to the membrane vesicular furrow initials via the centrosomal region and are then dragged inwards when the vesicular furrows are elongated. (A &. **B)** membrane vesicular furrows form in the proximity of the nuclear centrosomal regions (yellow arrows; 5-to-15min). The nuclei remain tethered to the membrane via the centrosomal region when the vesicular furrows extend inwards (15-to-25 min). Note that in (A) a nucleus forms a spindle and remains attached via the polar region of the spindle to the extending furrow. Main scale bars 5 µm, inset 2 µm. **(C)** A strain expressing a fluorescently tagged centrosomal component SAS6-mClover3 shows that the centrioles are proximal to the vesicular furrow initials (yellow arrows) and remain attached to the furrows during elongation. Membrane dyed with FM4-64. Main scale bars 5 µm, inset 1 µm. See also Figure S2, and Video S5, S6.

To further understand the relationship between furrow formation and nuclear positioning, we followed nuclei, centrosomes, and membrane through the later stages of cellularization. Consistent with centrosomes playing a major role in organizing chytrid cellularization, we found that during furrow elongation (Stage 2) the centrosomes appear to remain tethered to the leading front of the furrow initials, dragging their nucleus with them (**Figure 3C; Video S6**). Taken together, these results suggest a physical connection between nuclei, centriolar basal bodies, and vesicular furrows that persists throughout cellularization.

### Microtubules are important for geometric patterning but are not necessary for cellularization

Microtubule-dependent forces help move and position nuclei and spindles during cytokinesis in animals and yeast. ^27–31^ Likewise, during *Drosophila* cellularization, microtubules position nuclei in a monolayer under the plasma membrane in preparation for cellularization, ^3,32,33^ and carry plus end-directed motors that help power early furrow ingression. ^3,34,35^ We therefore wondered if microtubules play similar roles in chytrid furrow elongation. To test this hypothesis, we treated cellularizing sporangia expressing fluorescent tubulin with 2 µM Nocodazole. Unlike cells treated with the DMSO carrier alone, sporangia treated with Nocodazole showed rapid dissipation of all microtubule structures except for a bright dot at the centrosome (**Figure S2B**). Cells lacking microtubules formed cellularization polyhedra of uneven size, curved edges, and rounded vertices **(Figure 4A)**. Despite these patterning defects, Nocodazole-treated sporangia completed cellularization and produced daughter cells lacking cilia (**Figure S2**). In contrast, sporangia treated with DMSO alone assembled well-organized cellularization polyhedra **(Figure 4A)** and produced normal, ciliated daughter cells.

**Figure 4.**
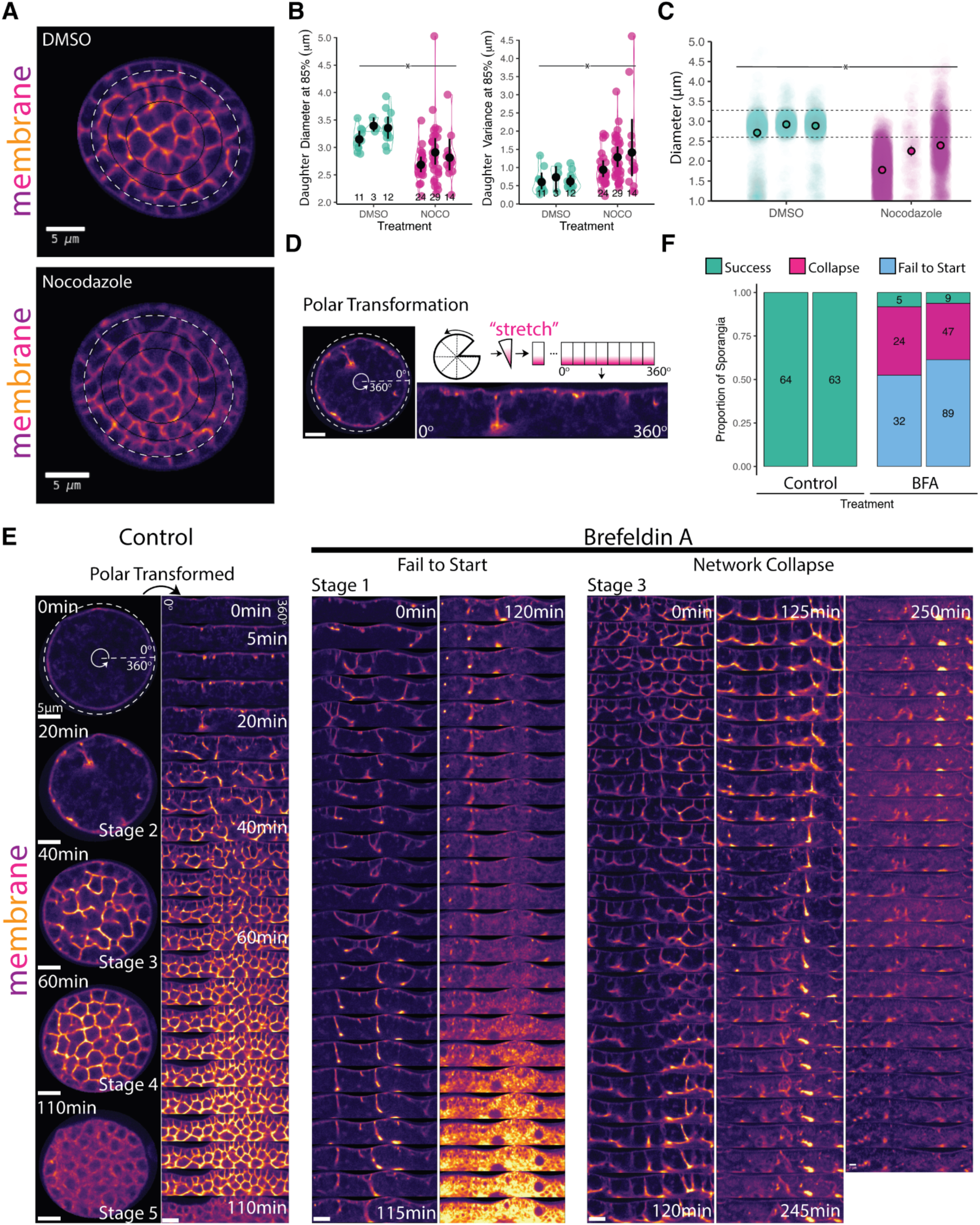
Depolymerization of microtubules does not inhibit cellularization but causes patterning defects and the membrane of cellularization is sourced primarily from the Golgi. **(A)** Sporangia treated with Nocodazole (2 µM) during cellularization causes a defect in the geometric patterning of cellularization but does not inhibit it. Concentric ellipses of 85% (white dashed line) and 65%, and 45% radii are shown in thin black lines **(B)** Daughter cells of sporangia treated with Nocodazole are smaller (left) and are more variable (right) in diameter than sporangia treated with the carrier control DMSO. The daughter size (daughter diameter) was estimated using the distance between max-intensity peaks in a line scan of the fluorescence intensity of the membrane along the 85% radii ellipse. Cellularization planes at 85% tend to be perpendicular to the ellipse and thus more accurate. Significance was assessed by a permutation test of the difference of means. N=3; * p-value ≤ 0.05. **(C)** The effect of Nocodazole on zoospore size was confirmed by treating synchronous cultures of *Spizellomyces* with 2 µM Nocodazole for two hours upon entry to cellularization and measuring the size of the zoospores produced using a Coulter Counter. We found that Nocodazole causes a significant reduction in average zoospore size (N=3, n=2412, 2227, 3644, 4466, 183, and 2186, permutation test of difference of means: * p-value ≤ 0.05). Although the variance appears to be affected, we found this difference not statistically significant (permutation test of difference of means, p-value=0.1). **(D)** Schematic of the polar transformation on an image of membrane organization (FM4-64) in a sporangium during cellularization. This transformation converts “circular” images with polar coordinates into “unwrapped” images in cartesian coordinates for easier visualization. In our case, the plasma membrane will be at the top and the furrows invade inwards towards the bottom. **(E)** Sporangia treated during cellularization with the inhibitor of ER-to-Golgi trafficking Brefeldin-A fail to complete cellularization because they either fail to start—furrows try to invade but fail to progress (left)—or cause collapse of the furrows already formed at the moment of treatment (right). In contrast, control sporangia treated with the carrier DMSO show a gradual increase in the complexity of patterning until completing cell separation (110 min). Bright FM4-64 fluorescence in the “fail to start” panel corresponds to cell death. **(F)** Quantification of the proportion of cells succeeding, failing to start, or collapsing during treatment with BFA or DMSO control. All scale bars are 5 µm. See also Figure S2.

To quantify the patterning defect induced by microtubule depolymerization, we compared the size and variance of the cellularization polyhedra between Nocodazole-treated and control cells. To aid automated image analysis, we focused on the polyhedra contiguous to the cell wall because they have regular, perpendicular furrows (**Figure 4A**). We found that, while sporangia treated with DMSO produced polyhedra with diameters between 3-3.5 µm, sporangia treated with Nocodazole produced polyhedra between 2.5-3 µm—a 14% reduction in diameter, with an even greater change in variance (**Figure 4B**). We confirmed this finding by measuring the diameters of fully developed ciliated daughters—zoospores—by treating synchronous cultures with 2 µM Nocodazole upon entry into cellularization and harvesting the zoospores released after two hours of treatment (**Figure 4C**). We found a consistent and significant reduction in daughter size relative to DMSO control cells (diff of means one-tail permutation test=0.6973, p-value 0.05). Microtubules, therefore, are not strictly necessary for cellularization but do play an important role in patterning the membrane geometry and regulating polyhedral/daughter size.

### The additional membrane needed for cellularization is sourced from the Golgi

Building the cellularization foam requires large amounts of membrane. Since the membrane used for cytokinesis in animals and yeasts comes from the Golgi, we wondered if this was also the case for chytrid cellularization. To test this hypothesis, we disrupted ER-to-Golgi trafficking with Brefeldin A (BFA). We treated synchronous populations of *Sp* with 50 µM BFA—or DMSO control—in the presence of FM4-64 and followed sporangial development by time-lapse imaging for six hours (**Figure 4D-F)**. To help visualize the cellularization pattern, we computationally “unwrapped” the circular sporangia by polar transformation (**Figure 4D)**. While control sporangia completed cellularization (N=127), only 7% of sporangia treated with BFA completed cellularization during the same period of time (N=206), 12 out of 14 of which had already started cellularization when BFA was added **(Figure 4E)**. The sporangia that failed to cellularize displayed two main phenotypes. The first was “failure to start”. Sporangia with this phenotype were in either Stage 1 or Stage 2 upon BFA treatment **(Figure 4E)**. The furrows of these cells made repeated attempts to elongate but never progressed. These cells eventually arrested cytoplasmic movement, vesiculated, and acquired intense, uniform FM4-64 fluorescence—suggestive of cell death. The second phenotype was “network collapse”, which occurred in sporangia that already had an advanced cellularization network (Stage 3) prior to BFA treatment (**Figure 4E**). The cellularization structures in these sporangia collapsed into a single disorganized bundle followed by a death similar to that in the “failure to start” phenotype. Together, these results show that ER-to-Golgi trafficking is necessary for cellularization. This result agrees with ultrastructural imaging of other chytrids, where cellularization coincides with the hypertrophy of Golgi cysternae and proximity of these structures to cellularization furrows. ^36^ Although we cannot reject contributions from additional endomembrane systems, our phenotypes suggests that the membrane for chytrid cellularization is sourced primarily from the Golgi.

### Actomyosin contraction is a driving force of chytrid ceullarization

Having shown that microtubules are not necessary for chytrid cellularization, we next asked whether cellularization requires dynamic actin structures. We began by treating cellularizing sporangia expressing a fluorescently-tagged F-actin probe (Lifeact) with Latrunculin B (LatB)—a reversible inhibitor of actin polymerization and accelerator of depolymerization. ^37,38^ Treatment with LatB caused disassembly of the actin cellularization networks and partial disassembly of the cellularization membranes within five minutes (**Figure 5A; Figure S4**). This effect was reversible, as the cellularization networks began reforming within five minutes of LatB wash out, indicating that these networks are undergoing dynamic turnover. We quantified these phenotypes using fluorescence intensity variability (*i.e.* Haralick image texture) as a proxy for the structural complexity of the actin networks (**Figure 5B**). While sporangia treated with ethanol alone showed no significant change in texture, sporangia treated with LatB showed a 50-75% increase in homogeneity within five minutes of treatment. Washout of LatB returned texture levels to pre-LatB levels within five minutes. Because Latrunculin B inhibits actin assembly, these results indicate that cellularization networks require actin polymerization.

**Figure 5.**
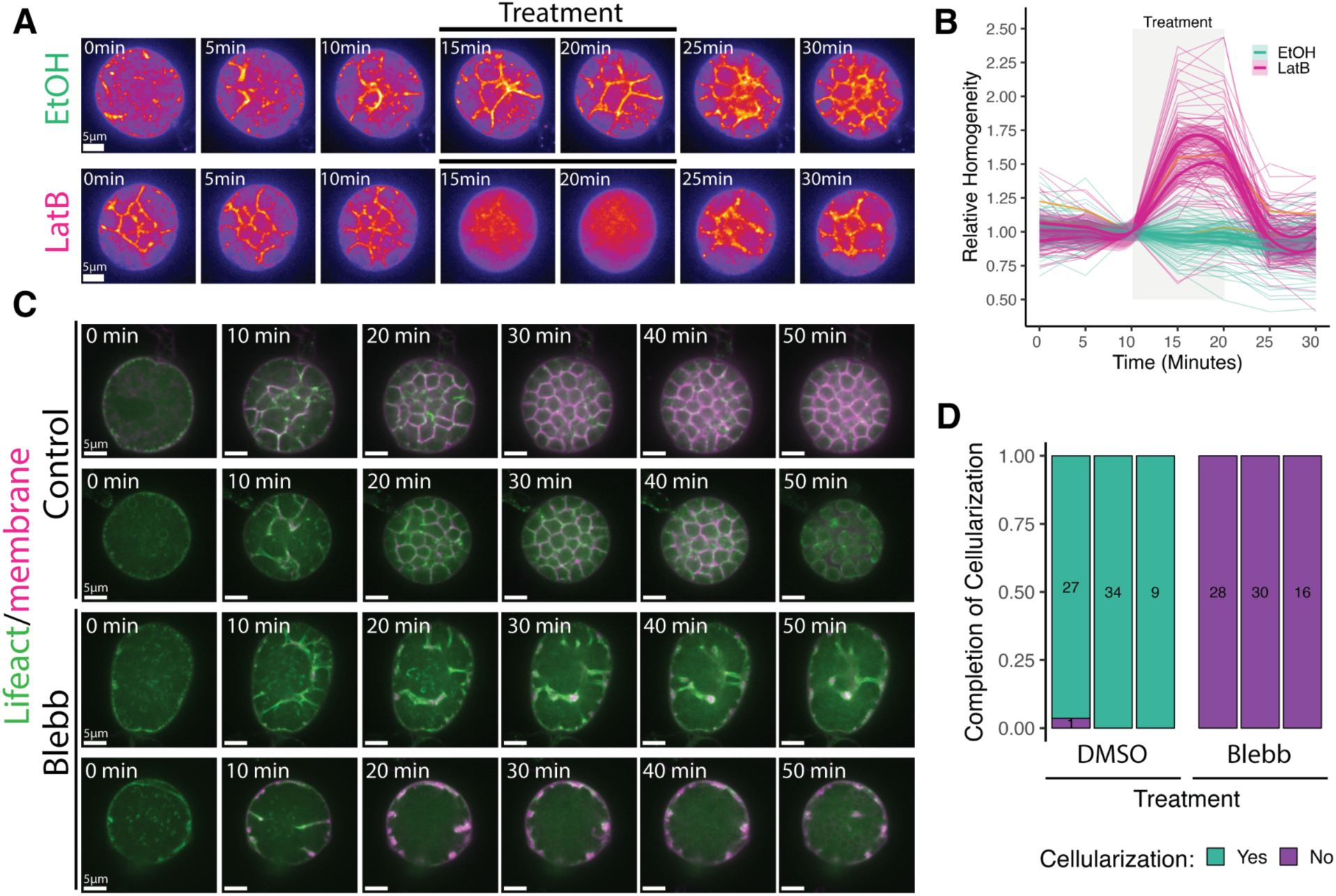
Chytrid cellularization requires actin polymerization and myosin-II activity. **(A)** Inhibition of actin polymerization with Latrunculin B (LatB) in a strain expressing the actin probe Lifeact-mClover3 causes disassembly of the actin cellularization network. This network is reassembled within 5 min after the washout of LatB. **(B)** Treatment with LatB (magenta) causes a sharp increase in sporangial homogeneity—reduction in texture, and thus disassembly of the actin network—that comes back to pre-treatment levels after the washout of LatB. Control sporangia (teal) treated with DMSO control shows no effect. The mean line and bootstrapping 95% confidence interval of the mean are shown for three biological replicates. Orange lines show the series shown in panel (A). **(C)** Treatment of sporangia with the inhibitor of myosin-II activity para-amino-blebbistatin (Blebb) inhibits cellularization. Control cells treated with DMSO control show normal cellularization, while sporangia treated with Blebb fail to advance beyond Stage 1 or 2. **(D)** Treatment of sporangia with Blebb completely inhibits cellularization. The box for each of the three biological replicates shows the number of sporangia analyzed per replicate. All scale bars are 5 µm. See also Figure S4.

In animals and yeast, similar-looking actin networks collaborate with myosin-II motors to generate the contractile forces that drive cytokinesis. To test if myosin-II is necessary for chytrid cellularization, we treated cellularizing sporangia expressing Lifeact with the myosin-II inhibitor para-amino-blebbistatin (blebbistatin). While sporangia treated with the carrier control DMSO underwent normal cellularization, sporangia treated with blebbistatin failed to complete cellularization (**Figure 5C & D**). These results show that both actin polymerization and myosin-II activity are required for chytrid cellularization.

The actomyosin networks that drive cytokinesis in animals and yeast generate tension that can be detected by ablating actomyosin structures in a diffraction-limited spot and observing their responding motions. ^39–44^ If chytrid cellularization is powered by a similar contractile mechanism, then severing the actin furrows should cause remaining structures to recoil as they release tension ^40,41,43,44^ We therefore used laser ablation to test if the actin networks of early and late cellularization are under tension and observed recoils in furrows of Stage 1 (**Video S7**) and Stage 3 sporangia (**Figure 6A; Video S7**). Ablation of Stage 3 furrows caused large recoils of actin structures that later moved back into the ablated area (**Figure 6A** kymographs), a “repair” response observed in other contractile actin networks. ^40,41,43,44^ To quantify the speed and direction of the large Stage 3 recoil, we turned to particle image velocimetry (PIV) (**Figure 6B-D**). By using pixels from the actin network images as tracer particles, we could account for the variability in response across cellularization structures in different sporangia. This analysis shows that, after laser ablation, many of the F-actin structures within a few µm of the damaged region move with high velocities away from the ablation site (**Figure 6B**), indicating that neighboring contractile structures likely distribute their load onto each other.

**Figure 6.**
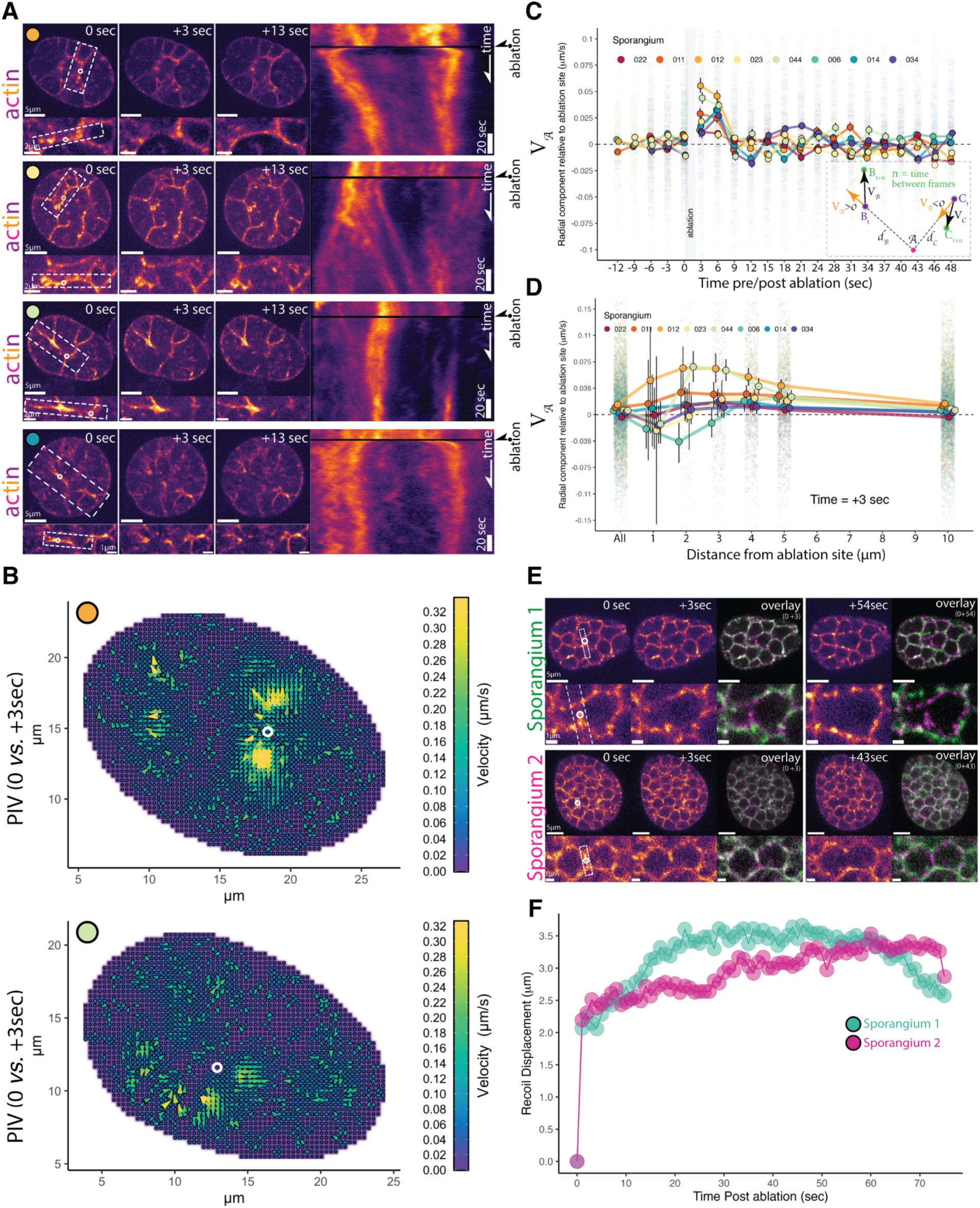
The actomyosin furrows of early cellularization and the edges of the cellularization foam are under tension. **(A)** Laser ablation of the actin furrows during early cellularization results in recoil. Four examples of sporangia ablation are shown. Colored circles correspond with sporangia shown in later panels. Each row shows still images of the sporangia just before laser ablation (0 sec, at white circle) and then 3 and 13 seconds after laser ablation, with dashed region inset below. The rectangle in each inset shows the 20-pixel wide selection used for the sum of intensity kymograph on the right. In each kymograph, the fast recoil response after the dark rows (ablation) is followed by contraction of the actin structure as if “repairing” the damaged region. Main scale bars 5 µm, inset scale bars 2 µm, 2 µm, and 1 µm, from top to bottom. **(B)** We used particle image velocimetry (PIV) to quantify the changes in the speed and direction of all actin structures of cellularization. Two examples (sporangia with corresponding color circles in panel A) of PIV velocity field plots comparing the frames before laser ablation (0 sec) and after ablation (3 sec). Note vectors with high velocity (warm color) moving away—recoiling—from the ablation sites (white circles). **(C)** Laser ablation causes a high velocity recoil away from the ablation site in all actin structures within 4 µm of the ablation site. We use the radial component of the velocity (V *_A_*) relative to the ablation site (*A*) to show movement only in the direction of the ablation site (away—postive—or towards—negative; see cartoon in inset). The short period of negative V *_A_* that follows the recoil in most sporangia is consistent with the “repairing” contraction towards the ablation site. Color dots show mean values (corresponding sporangia shown with the same color circles of A) and error bars show 99% confidence intervals of the mean by bootstrap. For visualization purposes we display data points within a range of -0.1 to 0.1 µm/s (this is 176417 points; 95% of the total data). **(D)** Radial component of the actin velocities (V *_A)_* relative to the ablation site (A) for vectors at different distances from the ablation site (*d* in cartoon) at three seconds after the start of ablation. For visualization purposes we display data points within a range of -0.15 to 0.15 µm/s (this is 23198 points; 97% of the total data). Note that highest velocities occur between 2-4 µm from the ablation site in most sporangia. **(E)** Laser ablation of the foam furrows during late cellularization also results in recoil and dilation of the polyhedral geometry. Overlay for frames before and the first frame after the end of ablation (3 sec from the “before” image) and between the frame before and the frame showing near maximal recoil displacement (54 sec & 43 sec). Main scale bars 5 µm, inset 1 µm. **(F)** Recoil displacement caused by ablation of the sporangia (pink and green) shown in E, along the direction indicated by the white dotted rectangle in the inset. The recoil of the polyhedral edges shows a sharp fast recoil of about ∼2 µm—primarily from recoil of the foam furrow—followed by dilation of the surrounding polyhedra. See also Figure S5, S6, and Video S7, S8.

To characterize the time scale and spatial extent of the response to ablation, we analyzed the radial component of the PIV motion, either away from (recoil) or toward (repair) the site of ablation at different distances from the targeted region (**Figure 6C, inset**). Consistent with the qualitative observations above, this analysis demonstrates an initial recoil response—indicated by a positive radial component of the velocity lasting around nine seconds—followed by a short period of movement towards the ablation site consistent with repair of the damaged structures—indicated by a negative radial component of the velocity (**Figure 6C**). While we see variation in timing and magnitude among different sporangia, this overall signal is robust to different strategies of pre-processing of the data (**Figure S5**). We found the fastest recoil signal occurred between two and four µm from the ablation site (**Figure 6D, Figure S6**) and between three and six seconds after ablation (**Figure 6C, Figure S6**). Due to the comparable timescales of recoil (∼1-9 seconds), the ablation (∼1-2 seconds), and our imaging frequency (1 Hz), there is significant uncertainty in the instantaneous magnitude of the recoil velocity, which may be faster than the ∼75 nm/s we can detect (**Figure 6C-D**). These results show that the actomyosin structures of early cellularization are contractile, under tension, and distribute their load over a distance of several µm.

Finally, we tested whether the F-actin polyhedra of late cellularization are also under tension. We found that ablation of the edges of the cellularization polyhedra resulted in a rapid recoil of the furrow tips away from the ablation site (**Figure 6E; Video S8**), showing a displacement of approximately two µm within the first second after ablation followed by a shallow exponential recoil profile in which tension appear to release for more than 30 seconds (**Figure 6F**). After the first recoil, the two vertices at each side of the ablated furrow also recoiled away from each other, causing a dilation in adjacent polyhedra. Together, these results show that contractile actomyosin is used during furrow elongation as well as throughout the assembly of the cellularization network, and is important for establishing and maintaining the geometry of the cellularization foam.

## Discussion

Here, we dissect the process of chytrid “volumetric” cellularization into molecularly distinct stages (**Figure 7**). We find that cellularization onset is marked by the migration of nuclei and their associated, outward-facing centrosomes to the plasma membrane, followed by formation of a vesicular invagination at the point where the centrosome abuts the plasma membrane. These vesicular structures then project inward in an actomyosin-dependent process, becoming tubular “furrows” that drag attached nuclei into the cytoplasm. The tubular furrows expand, branch, and merge using membrane sourced from the Golgi and eventually settle into a foam of polyhedra of homogenous volume, each containing a single nucleus. Finally, the polyhedra synchronously separate from each other to form daughter cells that crawl or swim away through a pore in the cell wall of the mother. During cellularization, one of the two centrioles associated with each nucleus transitions into a basal body and grows a cilium. We show that, although chytrid cellularization requires actomyosin networks, it uses them in ways that are drastically different from those of animals and yeast.

**Figure 7.**
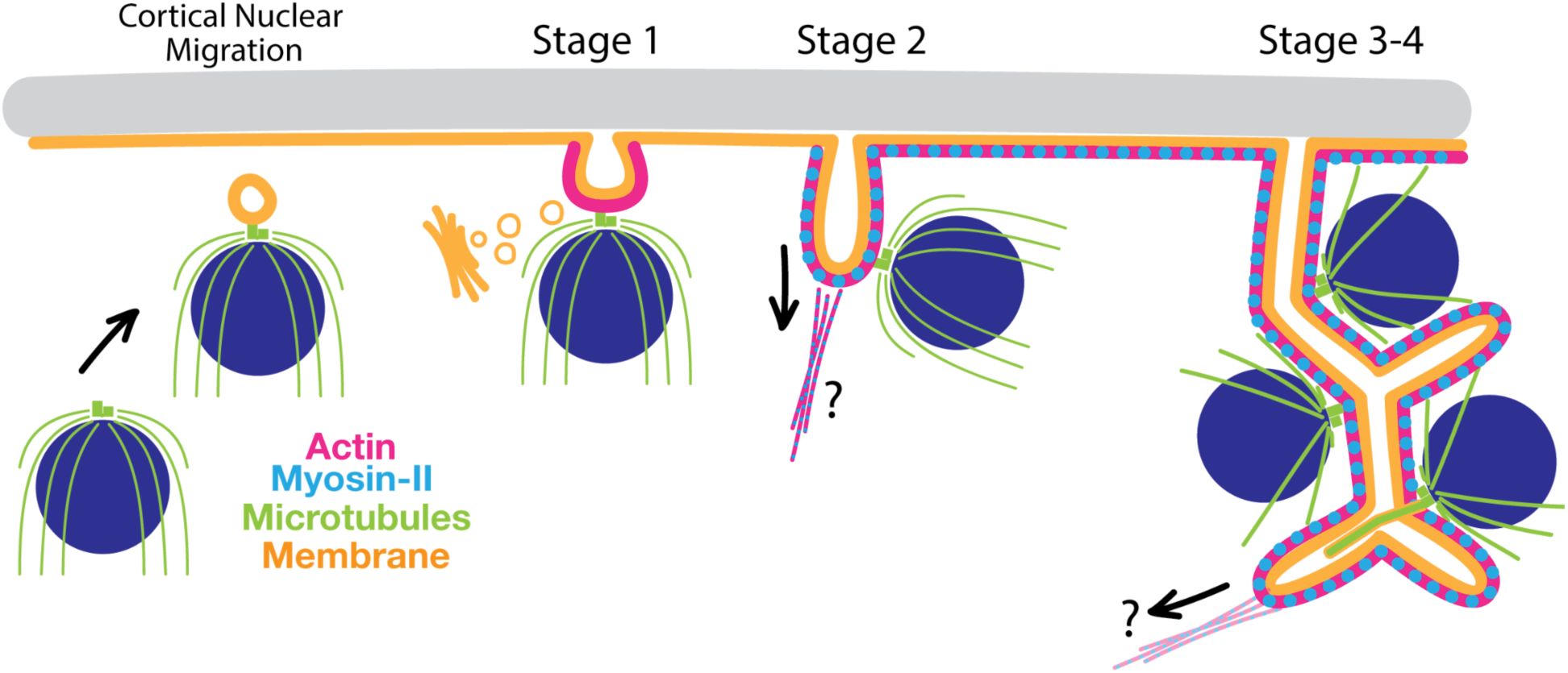
The model of chytrid aphrogenic cellularization. Nuclear cortical migration: Entry into cellularization is marked by the recruitment of nuclei and associated centrioles to the plasma membrane. We hypothesize that a “ciliary vesicle”-like structure may be associated with one centriole during nuclear migration. After migration, the centriolar centrosome of each nucleus faces the plasma membrane. **Stage 1, Vesicular furrow initiation:** the ciliary vesicle fuses to the plasma membrane—allowing the rapid diffusion of FM4-64 dye from the plasma membrane. This ciliary vesicle is also the vesicular furrow initial. This stage does not depend on myosin-II and the membrane sourced from this stage onwards is derived from the Golgi apparatus. **Stage 2, Furrow ingression:** Actin and myosin-II work together to drive the extension of the vesicular furrows into tubular furrows that invade the cytoplasm. Because the nuclei remain attached to their furrows via the centrosomal region, the extension inwards of the furrows drags the nuclei with them into the cytoplasm. We infer that nuclei dock during furrow initiation (Stage 1) because they move with the furrows during furrow ingression. The location and organization of the connections in the actomyosin network that allows for the “pulling” inwards of the furrows from the plasma membrane are yet unclear. Nuclei tethered to the furrows can be dragged inwards alone or along with additional nuclei, suggesting connections between nuclei may form prior to furrow invasion. **Stage 3-4, Aphrogenesis: assembly of the cellularization foam.** Tubular furrows, comprising membrane and associated actomyosin networks, branch and merge to generate the polyhedral territories of cellularization, each containing a single nucleus. The microtubules play a role in patterning the size and geometry of the polyhedral territories. The ciliary axonemes extend in between the leaves of the double-membrane of the polyhedral faces. How the tubular furrows generate the polyhedra and how tubular structures transform into sheets is unclear. **Stage 5, not shown:** Finally, after maturation of the cellularization foam, the zoospores undergo cell separation and swim or crawl out through an opening the the cell wall of the mother.

### Chytrids use actomyosin for cellularization but with different mechanisms from animal and yeast cytokinesis

We use the spatial arrangement of nuclei to categorize the geometry of cellularization as in-line (conventional cytokinesis), in-plane (e.g., *Drosophila* cellularization), or volumetric (chytrid fungi), and show that chytrids use actomyosin in unique ways to drive volumetric cellularization. Conventional cytokinesis and *Drosophila* cellularization rely on the assembly of an actin ring at the plasma membrane that constricts and deforms the membrane into a trench-like furrow around each nucleus. Similarly, chytrids use actomyosin to drive membrane invagination during cellularization. However, chytrid cellularization requires building furrows in regions of the cell that do not have ready access to the plasma membrane. Instead of building furrows around each of the internal nuclei, chytrids recruit nuclei to the plasma membrane prior to furrow initiation and tether each nucleus to a furrow. Although our data lack the resolution to determine if *all* nuclei must be recruited to the plasma membrane for proper cellularization, nuclei that do migrate and tether to the plasma membrane maintain their connection to the plasma membrane during furrow invasion. The resulting tubular furrows are then used to build the cellularization foam. Thus, chytrid cellularization neatly solves a problem not faced by well characterized animal or yeast model systems whose nuclei are in proximity to the plasma membrane.

Similar mechanisms to chytrid cellularization are likely deployed by other fungi that generate spores from a multinucleated sporangium, including two groups of emerging fungal pathogens, members of Mucoromycotina and *Coccidioides.* Mucoromycetes include *Mucor* and *Rhizopus* that cause the deadly black fungus disease that followed the COVID-19 pandemic. ^45^ These fungi produce spores by a mechanism of multinuclear cellularization that includes invasion of tubular membranes and polygonal territories around daughter nuclei. ^46–51^ Fungi of the genus *Coccidioides* cause the Valley fever respiratory disease and form a multinucleated cell—called a spherule—that cellularizes into a honeycomb of uninucleated polygonal endospores. ^52–55^ Chytrid multinuclear cellularization provides a framework to understand the developmental biology of these important human pathogens.

Actomyosin-based multinuclear cellularization is not limited to fungi and *Drosophila*. The Ichtyosporeans *Sphaeroforma arctica* and *Creolimax fragrantissima* are unicellular relatives of animals that, like *Drosophila*, undergo in-plane cellularization to produce an epithelia-like monolayer of cells. ^56–58^ Indeed, long term term treatment of *S. arctica* with a microtubule depolymerization agent (Methyl Benzimidazol-2-yl-carbamate; MBC) suggest that, like chytrids, Ichtyosporeans may use microtubules in nuclear positioning but not cellularization ^56^ The plasmodium of the acellular slime mold *Physarum polycephalum* also likely uses actomyosin for multinuclear cellularization. ^59,60^ Likewise, based on transitions from multinucleate to uninucleate cell states, multinuclear cellularization may also occur in slime mold sporocarps, structures that were likely present in the ancestor of the Amoebozoa. ^61^ Because the shared ancestor of animals, fungi, and slime molds likely had a multinuclear stage, ^62^ it may have had the capacity to generate uninucleated propagules or gametes through multinuclear cellularization. Here we show that, like animals and Ichtyosporeans, chytrid fungi rely on actomyosin networks to drive multinuclear cellularization, suggesting that using actomyosin for this function may be an ancestral feature of animals and fungi.

Although chytrid fungi are the first system—to our knowledge—shown to use actomyosin for volumetric cellularization, they are most certainly not the only system capable of “volumetric” cellularization. This strategy is present in many eukaryotic lineages, including land plants and Oomycetes. During land plant cell division, a specialized structure called the phragmoplast forms a cell plate in the center of the dividing cell through polarized delivery of membrane and cell wall components. This outgrowing disk eventually reaches the plasma membrane and partitions the cell cytoplasm. ^63^ Endosperm tissue of many plant species cellularizes by building mini-phragmoplasts between all of the nuclei. ^14,64,65^ More similar to chytrid cellularization, Oomycetes like the plant pathogen *Phytophthora* or the fish pathogen *Saprolegnia* also produce motile zoospores via cellularization of a multinucleated sporangium. ^66^ Like chytrids, cellularization in these species begins with the recruitment of Golgi-derived vesicles to each nucleus’ centrosome. Oomycete sporangia then form membranes around the cell periphery that extend and merge with the centrosome-associated vesicles to surround each individual nucleus.^13,67^ These differences in mechanism and strategy used in different lineages leave open the question of whether cellularization evolved multiple times or was ancestral and has undergone rampant diversification.

### Chytrid cellularization combines ciliogenesis and cytokinetic mechanisms in a single cellularization program

Chytrids solve the problem of surrounding non-peripheral nuclei with cellularization membranes by moving internal nuclei to the plasma membrane before redistributing nuclei to the cytoplasm. This mechanism begins with three steps: 1) recruitment of the centriole-nuclei to the plasma membrane, 2) the formation of vesicular invagination between each nuclear centrosome and the plasma membrane, and 3) the tethering of the nuclei to these vesicular furrows via the centrosome. These events resemble key stages of animal ciliogenesis, which starts with the migration of the centriole to the plasma membrane. During this migration, the distal end of one centriole recruits membrane to form a double-membrane sheet called the “ciliary vesicle”. This centriole then fuses to the plasma membrane via the ciliary vesicle and extends an axoneme to form a cilium.^68^ Chytrid vesicular furrow initials resemble animal ciliary vesicles not only in their appearance and position, but also in their genesis. Upon recruitment to the plasma membrane, chytrid nuclei interact with the vesicular furrow initials via the centrosomal region, an association that persists throughout the rest of cellularization. We therefore propose that nuclear tethering during chytrid cellularization may be equivalent to centriolar docking in animal ciliogenesis, which relies on genes associated with human ciliopathies ^69^. Many of these genes are also present in chytrids, but whether chytrids use them for ciliogenesis is an open question. Finally, ultrastructural studies of chytrid ciliogenesis show that it begins by forming a “primary flagellar vesicle” similar to the animal “ciliary vesicle”. ^20–22,70^ The “primary flagellar vesicle” can be seen during chytrid ciliogenesis at the distal part of the basal body and against the plasma membrane. We propose that this “primary flagellar vesicle” *is* the vesicular furrow initial. This would mean that ciliogenesis and cellularization in chytrids are intertwined, spatiotemporally and mechanistically.

The similarities between animal ciliogenesis and early chytrid cellularization extend to their cytoskeletal mechanisms. We found that Nocodazole treatment did not inhibit cellularization but did inhibit axonemal elongation. This is similar to ciliogenesis in the quail oviduct, where Nocodazole treatment did not inhibit centrosome migration or centriolar docking but did inhibit axoneme elongation. ^71,72^ Similarly, actin network perturbation disrupts cellularization in chytrids and centriolar migration in the quail oviduct. ^73^ Thus, chytrids may use ciliogenetic mechanisms ancestral to animals and fungi to recruit the centrosomes and their nuclei to the plasma membrane. Because mechanistic details of centrosomal migration during animal ciliogenesis remain unclear,^74^ cortical nuclear migration in chytrids is a fantastic model system to dissect the mechanisms of centrosomal migration and ciliogenesis in animals and fungi.

### Using chytrids to explore the physics and geometry of foams in biological systems

We have shown that chytrids cellularize by building a space-filling honeycomb. Although chytrid mother cells vary in size and number of nuclei, the daughter zoospores they produce are remarkably uniform in size. Daughter cell size is directly linked to the geometry of the cellularization polyhedra; polyhedra of homogenous volume give rise to daughters of homogenous volume. So how is the pattern, size, and symmetry of the honeycomb generated? A hint may come from its similarity to a system of packed soap bubbles, or “foam”. This soap bubble analogy has been invoked to explain foam-like patterns in biological systems that arise through distinct mechanisms, such as transmission of extracellular force across tissues, differential adhesion between cells, or space-constrained cell division. ^75–79^ These processes start from separate individual “bubbles” that interact to eventually give rise to the foam. Although chytrid cellularization resembles these types of foams, the genesis of the chytrid honeycomb is completely different. It does not arise from the agglomeration of separate, individual “bubbles”. Instead, all of the interfacing surfaces and individual polyhedra in the foam are constructed *de novo*. Because chytrid cellularization entails the generation of a foam, and names such as “3D cellularization” or “volumetric” cellularization fail to communicate the multidimensionality and geometry of this process, we have named this process *aphrogenic cellularization* (foam-producing; from the greek αφρóς = aphros = foam)

Putting chytrid cellularization in the context of a geometric foam explains some of the unusual geometry of aphrogenic cellularization. Generating zoospores is energetically expensive, and evolution may have selected for the most efficient way to compartmentalize a sporangium into zoospores of defined volume. Under this model, chytrids have evolved to solve a special case of Plateau’s problem called *Kelvin’s Problem—* finding the most efficient “tessellation of space into cells of equal volume with the least surface area”. ^80^ The solution proposed by Kelvin of a 14-sided truncated octahedron with eight slightly distorted hexagonal faces and six square faces remained unchallenged for more than 100 years until Weaire and Phelan used simulations to propose a structure that improved upon Kelvin’s by 0.3%. ^81^ The Weaire-Phelan foam is made up of two types of cells, a 12-sided irregular polyhedron of pentagonal faces and a 14-sided polyhedron with pentagonal and hexagonal faces, with only the hexagons remaining planar. Although we lack a detailed understanding of the exact geometry of chytrid cellularization foams, we found very few instances of right angles, suggesting a structure more akin to the Weaire-Phelan than Kelvin’s. Because Kelvin’s problem must be solved by all biological systems that involve complete cellularization within a constrained volume, understanding the mechanisms used by chytrids to solve Kelvin’s problem can shed light on the principles of cellular morphogenesis across eukaryotic lineages, as well as inspire new strategies for solving geometrical problems within or without biological systems.

The soap bubble analogy can also help us think about how chytrid mothers produce homogenous daughter cells. Our observations suggest that chytrid cellularization obeys at least the first three of Plateau’s Laws of foams. Whether chytrids comply with the fourth Law—that surfaces meet in fours at a tetrahedral angle—awaits high resolution volumetric imaging of the cellularization foam. If we assume that chytrid cellularization follows Plateau’s Laws, then chytrids cellularize under three spatial constraints: a fixed mother’s volume, all compartments have the same “target” volume, and each compartment must contain a single nucleus. This raises important questions regarding the molecular mechanisms underlying how the target cell volume is defined and how the mother builds a compartment around each nucleus—aphrogenesis. Given that microtubule depolymerization results in patterning defects, microtubules may help define proper nuclear spacing and a “target volume”. Additionally, tethering the nuclei to the cellularization furrows may help keeping track of the nuclei and serve as a guide around which to build each polyhedron.

### Concluding remarks

Chytrid cellularization solves three-dimensional problems that neither animal nor yeast model systems face. Accordingly, we have found that chytrids fulfill each of the four fundamental steps of cellularization using mechanisms that consistently deviate from the known strategies deployed by animal and yeast model systems. First, in animal and yeast cytokinesis, nuclei find and hold their position while cellularization furrows move around them. In contrast, chytrid nuclei are highly dynamic throughout cellularization. Second, animal and yeast systems rely on plasma membrane-associated cues to recruit and assemble the cytokinetic machinery. While conceptually simple for in-line and in-plane cellularization, this constraint is problematic for volumetric cellularization because most nuclei lie deep in the cytoplasm, surrounded by other nuclei, and are therefore spatially separated from plasma-membrane bound cues. Chytrids solve this problem by recruiting and attaching the nuclei to the plasma membrane, and *then* using actomyosin to build cellularization structures away from the plasma membrane. Third, while animals and yeasts use trench-like furrows comprising membranes and actomyosin as cellularization structures, chytrids use tubular furrows of membranes and actomyosin that only later assemble into a polyhedron around each nucleus. How the tubular structures of early cellularization become polyhedra and how these polyhedra separate into daughter cells remain open questions. Finally, we found that chytrid cellularization may have co-opted ciliogenesis programs, raising the question of whether the similarity between these forms of cell division reflects analogy or homology at the molecular level. Taken together, we have found that chytrids use conserved actomyosin machinery for aphrogenic cellularization but with different mechanisms from those of conventional cytokinesis and in-plane cellularization—a clear reminder that conservation of machinery does not always signify conservation of mechanism.

## RESOURCE AVAILABILITY

### Lead Contact

Requests for further information and resources should be directed to and will be fulfilled by the lead contact, Lillian Fritz-Laylin.

### Materials Availability

Plasmids generated in this study have been deposited to https://www.addgene.org/Lillian_Fritz-Laylin/

### Data and Code Availability

All data reported in this paper will be shared by the lead contact upon request.

All original code and analysis pipelines have been deposited at Zenodo and are publicly available at https://doi.org/10.5281/zenodo.17535393 as of the date of publication.

Any additional information required to reanalyze the data reported in this paper is available from the lead contact upon request.

## Acknowledgements

The authors wish to thank Iain Patten for valuable conversations, advice, and feedback and the NCSU Cellular and Molecular Imaging Facility. This work was funded by NIH grant 1R35GM138083 to M.W.E. and NIH grant 1R35GM143039 to L.K.F.-L. who is Canadian Institute for Advanced Research (CIFAR) fellow in the Fungal Kingdom: Threats and Opportunities program and an Investigator of the Howard Hughes Medical Institute. E.M.M. is a Howard Hughes Medical Institute Hanna H. Gray Fellow.

## Author Contributions

Conceptualization, E.M.M.; data curation, E.M.M.; formal analysis, E.M.M; funding acquisition, M.W.E., and L.K.F.-L; investigation: E.M.M.; methodology, E.M.M., M.W.E., and L.K.F.-L; project administration, M.W.E., and L.K.F.-L.; resources, E.M.M., M.W.E., and L.K.F.-L.; software, E.M.M.; supervision, M.W.E., and L.K.F.-L.; validation, E.M.M.; visualization, E.M.M., M.W.E., and L.K.F.-L.; writing –original draft, E.M.M., M.W.E., and L.K.F.-L.; writing – review & editing, E.M.M., M.W.E., and L.K.F.-L.

## Declaration of Interests

The authors declare no competing interests.

## Methods

### Experimental Model and Study Participant Details

#### Synchronized cultures for live cell imaging

Two days prior to the experiment, use Dilute Salts (DS) solution to harvest zoospores from an active culture of *Spizellomyces punctatus* (Koch type isolate NG-3) Barr (ATCC 48900) growing in K1 plates (1L; 0.6 g peptone, 0.4 g yeast extract, 1.2 g glucose, 15 g agar) supplemented with antibiotics (50mg/L Carbenicillin and tetracycline) and incubated at 30 Celsius with or without selection (200mg/L Hygromycin). ^17^ Upon zoospore release, filter zoospores through sterile 40-µm mesh filter (fisherbrand Cat. No. 22363547) followed by a sterile Whatman No1 syringe filter (∼11 µm). Plate 1mL of 1/5-1/10 dilution in DS solution of the zoospores into new K1 plates (right out of the fridge), seal with parafilm, and incubate for ∼20-24h at 28 Celsius until peak zoospore release—zoospores are ready for seeding the imaging plates. Dilute Salts solution (DS) corresponds to a 10X dilution of Machils medium B. ^82^ To prepare DS solution, make and filter-sterilize in advance two 2000X stock solutions (DS-A and DS-B) and aliquot into milliQ water when needed (500 µL each per Liter of DS). Recipe for DS-A and DS-B are in g/L. DS-A (KH2PO4:136 g/L, K2HPO4:174.2, (NH4)2HPO4:132). DS-B(MgCl2:9.52,CaCl2:11). To prepare the imaging plates, a cellvis 6-well glass bottom black plate with lid (P06-1.5H-N) is cleaned with a plasma cleaner for four minutes at maximum level (Harrick Plasma Expanded Plasma Cleaner 115V). Add promptly 1mL per well of 25ug/mL filter-sterilized 50% w/v polyethyleneimine (PEI) (sigma-Aldrich P3143-100ML) and incubate at room temperature for 10 min. After the 10 min, remove the PEI solution and rinse 10 times with 2mL of sterile DS solution, then add 2mL of sterile DS in preparation for inoculation. Do not let the treated surface dry out. Proceed to prepare the zoospore inoculum.

To seed the imaging plate, harvest and filter the zoospores from the semi-synchronous plate culture made the day before using the same approach as above. Quantify zoospore concentration with a hemocytometer by counting 10uL from a 30uL 1:10 zoospore dilution including 3uL of Lugol solution (Sigma-Aldrich; 62650-100ML-F) (killing zoospores and increasing contrast). Dilute the zoospores with sterile DS to achieve a working stock of 1×10^6^ zoospores per mL. Add 1mL of 1×10^6^ zoospores per mL of zoospore solution to each well and leave for 10 min without moving at room temperature. After 10 minutes, tilt and remove the excess zoospore solution and replace it with 2mL of pre-warmed 28C K1 liquid media (50mg/mL Carbenicillin; Fisher AAJ6194906), place on top of a 28C pre-warmed aluminum heatblock (two small tube heatblocks taped together) and leave for 10 min. After 10 min, wash gently three times with 2mL of liquid K1 (50mg/mL Carbenicillin), then fill with 3mL of liquid K1 (50mg/mL Carbenicillin), seal the plate with Parafilm and incubate inside a closed plastic container with a humidity chamber (empty pipette tip box with perforated tip rack filled with 50mL water). Incubate for ∼18h, at which point they are ready to start cellularization.

#### Construct and plasmid design

Constructs for this manuscript were synthesized by Twist Bioscience (South San Francisco, United States) and cloned by Twist directly into the binary *Agrobacterium* plasmid pCAMBIA1300 (Addgene: 44183). Plasmids were transformed into *Spizellomyces* by *Agrobacterium-*mediated transformation and transformant strains were recovered as described previously (see below). ^17,83^ *Synther8* is a short synthetic terminator (No 8) from. ^84^ Nuclear-localized mClover3 (GI3EM48; See **Table S1**) was generated using previously published pGI3EM22C ^17^ with Lifeact-tdTomato replaced by mClover3 with an N-terminal nuclear localization signal from *Spizellomyces* Rb protein SPPG_07796. Nuclear dynamics from Figure 2 were done with a strain carrying H2B-tdTomato from plasmid pGI3EM20C as previously described ^17^. We sequence verified all constructs by whole plasmid sequencing (Plasmidsaurus Inc.). Complete plasmid sequences and plasmids have also been deposited at Addgene (https://www.addgene.org/Lillian_Fritz-Laylin/).

## Method details

### Agrobacterium-mediated transformation

*Agrobacterium-*mediated transformation was performed as previously described. ^17^ *Agrobacterium* Strain EHA105 was used for all transformations. Plasmids were transformed via electroporation into competent *Agrobacterium* using 0.2 cm cuvettes in a Gene Pulser electroporator (Bio-Rad, USA) at 25 µF, 200 Ω, 2.5kV. Single colonies were streaked on selective plates (50 mg/mL; Kanamycin, Thermo Scientific J6127209). For chytrid transformation, a colony of transformed *Agrobacterium* containing the binary plasmid of interest was grown overnight at 30°C in 5 mL of Luria-Bertani broth supplemented with Kanamycin (50 mg/L). After centrifugation, the cell pellet was resuspended in 5 mL of Liquid Induction Media (IM-L), diluted to an OD660 of 0.1 in IM-L, and grown under agitation at 30C until achieving a final OD660 of 0.6, at which point the culture is ready for co-culturing with the chytrid (300 µL per transformation). Liquid induction media (IM-L) is composed of MM salts, 40 mM 2-(N-morpholino)-ethanesulfonic acid (MES), pH 5.3, 10 mM Glucose, 0.5% (w/v) Glycerol, and 100 µM Acetosyringone. ^17,85,86^ To prepare 200 mL of IM-L add 0.36 g of Glucose and 1 mL of Glycerol to 80 mL of 2.5X Minimal Medium Salts (K2HPO4 (5.125 g/L), KH2PO4 (3.625), NaCl (0.375), MgSO4.7H2O (1.25), CaCl2.6H2O (0.25), FeSO4.7H2O (0.00625), (NH4)2SO4 (1.25), there will be some precipitation after addition of CaCl2.6H2O, do not worry. Store at room temperature, no need to sterilize) and water to a volume of 190 mL. Autoclave the MMS with the Glucose and Glycerol and let cool down to approximately 50 Celsius. Dissolve 1.54 g of MES in 10 mL of MilliQ water and bring pH to 5.3 with KOH—the MES will dissolve better as pH approaches 5.3. Add 4 mg (for 100 µM) of Acetosyringone only after the pH has reached 5.3. Filter-sterilze and add directly to the rest of the autoclaved media (MMS, Glucose and Glycerol). Induction Media plates are made the same way, but recipe includes 20 g/L (2% wv) Agar and uses a final concentration of Acetosyringone of 200 µM (39 mg/L).

In parallel, harvest zoospores as described above from fresh *Spizellomyces* Wild Type cultures grown for at least three generations in K1 plates without antibiotics. Pellet the zoospores by centrifugation at 800 g for 10 min. We have found that one petri plate of active culture provides enough zoospores for a single transformation (one plasmid). Resuspend very gently the zoospores of a single plate in 300 µL of IM-L. For every transformation, zoospores and *Agrobacterium* are combined at four different ratios: 1:1, 1:0.25, 0.25:1, and 0.25:0.25 in a total volume of 200µL. To guarantee tight contact between *A. tumefaciens-S. punctatus* cells, the surface of the IM plate was rubbed with the bottom of a sterile glass culture tube to generate slightly concave depressions in each quadrant of the plate, wherein each 200µL co-incubation mixture was spotted. Plates were incubated unsealed for 4 days at room temperature. Mock transformations with empty *Agrobacterium* (no binary plasmid; grown in the absence of plasmid selective medium) were included as a negative control. After co-incubation, we added 1 mL of DS solution and gently scraped the plate with a razor blade, pooling the different cell ratios into a single 50 mL centrifuge tube, raising the volume to 20 mL with DS solution, and re-suspending clumps by inversion. The mixture was centrifuged at 1000 g for 10 min and the liquid phase was discarded. The remaining pellet was carefully resuspended with DS solution, plated on K1 plates containing Carbenicillin (50 mg/L) and Tetracycline (50 mg/L) to select against *Agrobacterium* and Hygromycin (200 mg/L) to select for transformed *Spizellomyces Spizellomyces* survival controls were performed by plating transformations after co-culture in non-selective K1 media (Carbenicillin (50 mg/L), Tetracycline (50 mg/L)). Transformation plates and controls were incubated at 30°C until colonies were observed (5–6 days). All plates were sealed with parafilm to prevent desiccation. Single colony isolates were retrieved by successive (five or six times) single-colony picking with a sterile needle, resuspended in DS solution and re-plated on a selective Hygromycin plate. For a video guide of the transformation procedure see ^83^

### Live cell microscopy and laser ablation

For live cell imaging, cells were imaged on a Nikon Ti2 microscope equipped with a Crest X-Light V2 L-FOV spinning disk (50 µm pinhole), a Prime95B sCMOS camera (Photometrics), and a Plan Apo λ 100x 1.45 NA oil objective, and an Okolab stage top chamber (H301-K-FRAME) with humidity control. The illumination setup was a Celesta Light Engine solid-state laser launch (Lumencor), Celesta VCGRnIR pentaband emitter 10-10857, and Celesta VCGRnIR penta-band dichroic 10-10858. mClover3 Emission filter ET535/70m (Chroma), FM4-64 and tdTomato emission filter ET610/75m emission. The microscope was controlled through NIS Elements software (Nikon).

Data for figures 1A and C, imaging plates were prepared as described above, but before imaging a small 1.5% low-melt agarose (Sigma, A9045) pad made in K1 media was placed on top of the cells. Images were acquired using a Leica Stellaris 8 STED mounted on a Leica DMI8-CS base and using an HC PL APO CS2 86x/1.20 water objective, and Power HyD spectral detector. Images were acquired with White Light Laser (WLL) at 0.85 power percentage, taken at 1.31X zoom, 16x line averaging, 1024×1024 resolution, and mClover target wavelengths of 509.62-586.36nm and FM4-64 target wavelengths of 624.50-771.73nm. For three-dimensional reconstruction in Figure 1A, images were acquired at 1024×1024×130 with voxel size of 0.0254x 0.0254 x 0.1484 µm and visualized in ChimeraX-1.61 ^87,88^ with gaussian smoothing of nuclei and membrane (sdev 0.05).

For laser ablation of F-actin structures during cellularization (Figure 6), imaging plates of a strain expressing Lifeact-mClover3 were prepared as described for live cell imaging but with a small (2mL) pad of 10% gelatin (Fisher Scientific G7-500) made in K1 media placed on top of the center of the well just before imaging. Imaging was done in a Nikon Ti-E microscope equipped with an Andor Dragonfly spinning disk confocal fluorescence microscope,100x NA 1.45 objective (Nikon), and a built-in 1.5x magnifier; 488 nm diode laser with Borealis attachment (Andor); emission filter Chroma ET525/50m; and an EMCCD camera (iXon3, Andor Technology).

Ablations were performed using a MicroPoint (Andor) system with galvo-controlled steering to deliver 20-30 3ns pulses of 551 nm light at 16-20 Hz (Andor) mounted on the Dragonfly microscope described above, as previously described. ^40,89^ Fusion software (Andor) was used to control acquisition while IQ software (Andor) was used simultaneously to control laser ablation. At this pulse rate, the ablation process lasts ~2 s. Chroma ET610LP mounted in the dichroic position of a Nikon filter turret was used to deliver the ablation laser to the sample. Since this filter also reflects the mEGFP emission, the camera frames collected during the ablation process are blank. The behavior of the severed F-actin structures of cellularization was imaged before and immediately following laser ablation by acquiring a single confocal plane in the 488-nm channel every second for at least one minute. The mechanism of ablative photodecomposition remains unclear but may be caused by either the propagation of a pressure wave and/or cavitation bubble dynamics. ^90,91^ The size of the damage is approximately the size of the diffraction spot of the lens <0.4 µm in the XY plane and <0.8 µm in the Z axis. ^92^

### Drug perturbations during live-cell imaging

About one hour before the start of the experiment (∼17h), remove excess of dislodged sporangia by removing gently 2mL of the media and replacing it with 2mL of fresh prewarmed K1+Carb media, adding and removing liquid from opposite sides of the well. Repeat this wash thrice in total. To the remaining 1mL, add 1mL of 20uM FM4-64 (stock 1mg/mL in water) made in K1+Carb media for staining the plasma membrane. Bring to the microscope and place in a preheated Okolab stage top chamber (H301-K-FRAME) with humidity control (30 Celsius). Twenty individual sporangia were selected from a single well for time-lapse imaging. Imaging started at 18h, sampling every 3 or 5 min, for six hours (for high-frequency sampling without pharmacological perturbations imaging was every 30s; See Video S6). Pharmacological perturbations are applied by removing 1mL of media and replacing it with 2X working solution of the drug of choice. For washout experiments, 50% of the final volume of the well is replaced with fresh pre-heated media ten times. Cells were treated with 2uM final concentration of Nocodazole (Cayman Chemical; No. 13857) or equivalent volume of DMSO, 50uM of Brefeldin A (Sigma B7651) or equivalent volume of DMSO, 10uM Latrunculin B (Millipore, 428020) or equivalent volume of ethanol, 150uM p-a-blebbistatin (Cayman 22699) or equivalent volume of DMSO.

### Coulter Counter cell size measurements

1mL of zoospores from six independent culture plates of *Spizellomyces* were freshly harvested as described in the section on semi-synchronized cultures for live imaging and were placed in a 25mL Accuvette (A35473; Beckman Coulter) with 24mL of freshly made 50% dilution of Isoton II diluent (C96980; Beckman Coulter) and run through a 10uM aperture tube (B42812; Beckman Coulter) on a Coulter Counter Multisizer 4e (B23005; Beckman Coulter) running on a freshly made 50% dilution of Isoton II diluent (C96980; Beckman Coulter). The Coulter counter is calibrated with 2-µm latex beads (C72513 CC; Beckman Coulter). The size distribution of zoospores and beads was exported as CSV, then processed, and visualized using JupyterLab 1.16 and R v4.1.2. ^93,94^

### Image processing and analyses

To measure the angle distribution in the polyhedral tessellation of cellularization (Figure 1E), five sporangia from three independent biological replicates of control cells stained with FM4-64 and imaged during cellularization were selected. The frame before obvious signs of cellularization was selected for each sporangium and the angles between the three cellularization planes (tri-furrow) around at least ten different nodes (where furrows meet) were measured using the angle function of Fiji (ImageJ2 v2.16.01.54p). ^95^ As a positive control and a gauge of technical reproducibility, artificial 120-degree tri-furrows with similar proportions of thickness-length were used for generating an artificial dataset by transposition and measured the same way at the same time. The resulting dataset was processed and visualized using JupyterLab 1.16 and R v4.1.2 facilitated by custom scripts. ^93,94^

For measuring nuclear cortical migration dynamics (Figure 2B) we used scikit-image ^96^ image processing in Python (3.9.16) in a Google Colab notebook. Briefly, elliptical sporangia were masked, and segmented and an ellipse was fitted to their perimeter. A second ellipse with 70% of radii from the perimeter ellipse was used as the boundary between the outer and inner 50% of the sporangial area. The nuclei were segmented via thresholding and watershed algorithms. The cortical nuclear ratio is the number of nuclei in the outer donut relative to the inner disk for each frame in the time-lapse video. The resulting measures were then processed and visualized using JupyterLab 1.16 and R v4.1.2 facilitated by custom scripts. ^93,94^

To measure the defect in patterning observed after Nocodazole treatment in cellularizing sporangia (Figure 4A & B), an ellipse was masked and fitted to circular sporangia as described above using scikit-image ^96^ image processing in Python (3.9.16) with custom scripts in a Google Colab notebook. Briefly, since the cellularization furrows of the outer layer of daughter cells are perpendicular to the perimeter cell wall, and thus size estimates are likely to be more accurate, a concentric ellipse at 0.85% of the fitted perimeter ellipse was drawn. The peak-to-peak distance along the profile of the FM4-64 fluorescence intensity along the perimeter of this ellipse was used as a proxy for daughter cell diameter. The resulting measures were then processed and visualized using JupyterLab 1.16 and R v4.1.2. ^93,94^ The statistical significance test was a one-tailed permutation test of the difference of means.

The effects of Brefeldin A on cellularization were visualized by similar sporangial segmentation and masking followed by log-polar transform using sci-kit-image ^96^ image processing in Python (3.9.16) with custom scripts in a Google Colab notebook. The phenotyping of the sporangia (Success, Failure to Start, Network Collapse) was done by hand in Fiji. ^95^ The resulting measures were then processed and visualized using JupyterLab 1.16 and R v4.1.2. ^93,94^

To measure the effects of Latrunculin B on the F-actin structures during cellularization, sporangia of circular shape that did not move excessively during the experiment were selected and imported to CellProfiler 4.2.1 ^97^ for segmentation. As a proxy of the level of cellularization in the actin structures, we used Haralick measures of texture, ^98^ the Angular Second Moment relative to the time-point right before LatB treatment—relative homogeneity—was used for visualization. The phenotyping of the effects of p-a-Blebbistatin on cellularization (completion or failure) was done by hand in Fiji. ^95^

For Particle Image Velocimetry (PIV) analyses, selected sporangia were masked in Fiji ^95^ and analyzed with OpenPIV-0.25.0 ^99^ through custom Python (3.9.16) in a Google Colab Notebook. Vector field plots (Figure 6B) and plots from the top 10% velocity vectors per PIV timepoint (Figure 6C) were generated in JupyterLab 1.16 and R v4.1.2 facilitated by custom scripts. ^93,94^ Kymographs from Figure 6A were generated using Fiji ^95^ from regions 20-pixel wide centered and parallel to the ablated furrow. Recoil displacement analyses from laser ablation of late-stage polyhedra were performed via sci-kit-image ^96^ image processing in Python (3.9.16) in a Google Colab notebook. Briefly, a 10-pixel wide region centered in a line along the ablated furrow was used to measure the mean fluorescence intensity profile along the furrow. Intensity peaks were detected and used to locate and measure the inter-peak distance between the receding tips of the ablated furrow. The results were visualized using JupyterLab 1.16 and R v4.1.2.^93,94^

A main concern during our analysis was the difference in time between image acquisition before and after ablation and the time required for ablation. PIV uses the displacement of tracer particles (pixels showing fluorescence of F-actin in our case) between consecutive images to estimate the changes in direction and velocity of the tracer particles. Although we imaged every second before and after laser ablation, it takes about two seconds to perform the 20-30 pulses of the laser required to sever the actin structures and one more second to remove filters and take the next image. Thus, our images before and after ablation are separated by three seconds, while in reality, there is about one second from the end of the laser ablation until the acquisition of the first image “post” ablation. Furthermore, we do not have the “real” image of the sporangium just before ablation, only before ablation is started. To navigate this problem we could assume that the sporangium before the start of ablation is a good proxy of the status of the actin structures before the end of ablation, and thus run our PIV analysis using a sampling time of one second before and after ablation. Alternatively, we can run the PIV analysis using three seconds as the sampling time between images, which would underestimate by up to three-fold the velocities in response to ablation, but it would be accurate about the true time difference between images. A second concern is that when all pixels within sporangium are used as tracer particles, we would detect displacement not only in cellularization structures but also movements of the fluorescent probe in the cytoplasm.

We tested different strategies of resampling and filtering the ablation data of a sporangium to assess their effects on the recoil signal retrieved by PIV (**Figure S5 & S6)**. We performed basic thresholding (MultiOtsu) to remove the cytoplasmic signal, resampled the complete time-series every three seconds to reflect the true time difference between pre-ablation and post-ablation images, combined thresholding and resampling, and combined resampling with a mask (generated from the sum of all frames before ablation followed by MultiOtsu thresholding) to restrict the PIV to the cellularization structures. To measure the displacement caused by ablation, we retrieved vectors with magnitude greater than zero and that pass PIV signal to noise ratio test and calculated their radial component of the velocity (V) relative to the ablation site (*A*) (**Figure 6C Inset**). This measure (V*_A_*) represents, for a velocity vector anywhere in the sporangium, the part of the velocity vector that is in the direction of the ablation site *A* This allows us to probe specifically the displacement velocities away (V *_A_* >0) or towards (V*_A_*<0) the ablation site irrespective of the complexity of geometry and movement of the actin structures across the sporangium. We found that the recoil signal is robust to different strategies of data processing, thus we decided to use three-second resampling of the dataset without thresholding as the main data processing for our analyses and discussion.

Quantification of recoil displacement after ablation of the structures of late cellularization (**Figure 6E-F**) were performed by calculating the distance between severed furrow tips using custom Python (3.9.16) scripts in a Google Colab Notebook and visualized and processed with JupyterLab 1.16 and R v4.1.2. ^93,94^. Tip position defined as the peak in the fluorescence intensity distribution in a line 10-pixel wide along the ablated cellularization plane (white dotted rectangle in inset).

### Quantification and Statistical Analysis

Custom Python scripts associated with the analyses in manuscript can be found on Zenodo (https://doi.org/10.5281/zenodo.17535393). All statistical values from the below analyses can be found in the main text and figure legends of this article. Statistical significance was assessed via permutation tests of the difference of means performed using Python 3.9.16 in a Google Colab Notebook (see image processing and analysis). Plots were generated with Python or with R v4.1.2 in JupyterLab 1.16. ^93,94^

## Supplemental Information

### Document S1. Figures S1–S6, Table S1 and supplemental references

**Video S1. Distribution of nuclei during chytrid stage 4 peak cellularization, related to Figure 1**. Nuclei (RbNLS-mClover3) shown in green and membrane (FM4-64) in magenta. Laser-scanning confocal imaging. Mother cell is about 20 µmmicrons deep: Z-plane sweep of 130 Z-stacks, each 0.148 µm. Scale 5 µm.

**Video S2. F-actin colocalizes with the cellularization membrane polyehedra of a live chytrid mother cell during stage 4 about to reach peak cellularization, related to Figure 1**. F-actin (Lifeact-mClover3) shown in green and membrane (FM4-64) in magenta. Note a small section is not completely cellularized. Laser-scanning confocal imaging. Mother cell is about 20 µm deep: Z-plane sweep of 138 Z-stacks, each 0.148 µmmicrons. Scale 5 µm.

**Video S3. Chytrid cellularize by assembling actin-based structures with complex geometry and architecture, related to Figure 1**. Three-dimensional visualization with ChimeraX of F-actin (Lifeact-mClover3; shown in green) during Stage 2 cellularization. Laser-scanning confocal imaging. Same data from Figure 1D.

**Video S4. Nuclear dynamics during chytrid cellularization, related to Figure 2**. Nuclei visualized with fluorescently-tagged histone H2B (H2B-tdTomato) are shown in green and membrane (FM4-64) in magenta. Note cortical nuclear migration at ∼298 minutes (left video) and ∼135 min (right video). Spinning-disk confocal time-lapse. Sampling rate of one image per minute. You can see the last two nuclear divisions before the chytrid mother enters cellularization. Scale 5 µm.

**Video S5. Microtubular dynamics and outward-facing nuclear centrosomes during entry into cellularization, related to Figure 3**. Fluorescently-tagged tubulin (mClover3-alpha-Tubulin) shown in green and membrane (FM4-64) in magenta. Mitotic spindles (5 min; Left) and centrosomal duplication (40 min; Left) are visible before entry into cellularization. During cortical nuclear recruitment, the nuclear-associated centrosomes are often positioned towards the plasma membrane (100 min in right video; 135 min in left video). Nuclei remain tethered to the invading furrow via the centrosome (140 min left video). Spinning-disk confocal time-lapse. Sampling rate of one image every five minutes. Scale 5 µm.

**Video S6. Centrosomal dynamics during chytrid cellularization, related to Figure 3**. Fluorescently-tagged component of the centriole SAS6 (SAS6-mClover3) shown in green and membrane (FM4-64) in magenta. Left video: Recruitment of the centrosomes to the plasma membrane precedes the formation of the vesicular furrow initials. Center video: The centrosomes remain tethered to the furrows during cellularization. Right video: complete cellularization process. Scale 5 µm.

**Video S7. The actin-based furrows of very early cellularization are under tension and recoil where severed by laser ablation, related to Figure 6**. Spinning-disk confocal time-lapse imaging of central plane of a strain expressing the F-actin probe Lifeact-mClover. The actin structures are severed using a laser. Note the fast recoil response after ablation. Scale 5 µm.

**Video S8. The actin-based furrows of early cellularization and the polyhedra of late cellularization are under tension and recoil where severed by laser ablation, related to Figure 6**. Spinning-disk confocal time-lapse imaging of central plane of a strain expressing the F-actin probe Lifeact-mClover. The actin structures are severed using a laser. Note the fast recoil response after ablation. Scale 5 µm.

## Notes

### Competing Interest Statement

The authors have declared no competing interest.

### Summary of Updates

The text has been revised throughout.

