## Supplemental Data for "Cellularization in chytrid fungi uses distinct mechanisms from conventional cytokinesis and cellularization in animals and yeast"

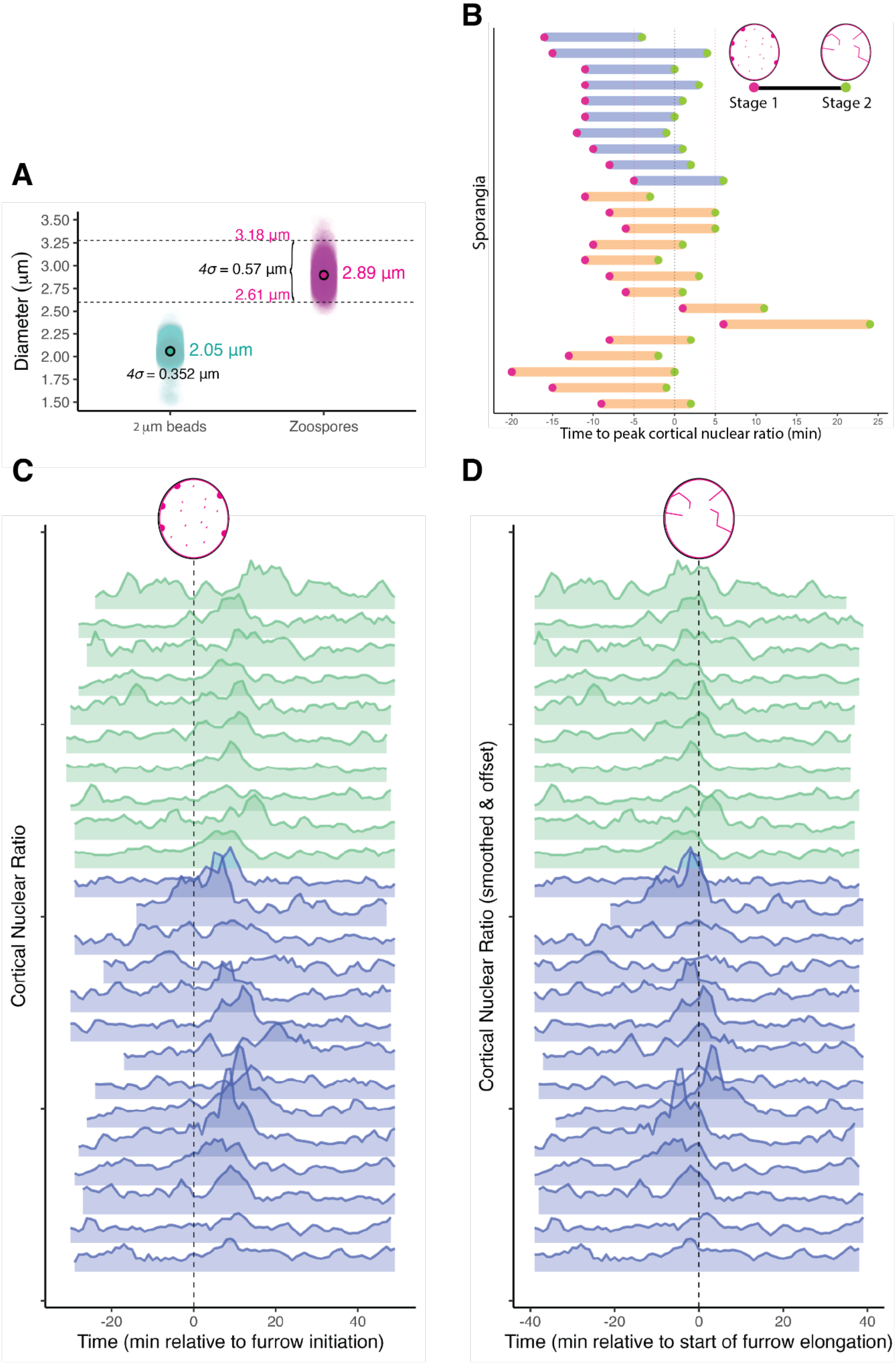

A
Diameter (µm)
3.50
3.25
3.00
2.75
2.50
2.25
2.00
1.75
1.50
3.18 µm
2.61 µm
4σ = 0.57 µm
2.89 µm
2.05 µm
4σ = 0.352 µm
2 µm beads
Zoospores
B
Sporangia
Stage 1
Stage 2
-20
-15
-10
-5
0
5
10
15
20
25
Time to peak cortical nuclear ratio (min)
C
Cortical Nuclear Ratio
-20
0
20
40
Time (min relative to furrow initiation)
D
Cortical Nuclear Ratio (smoothed & offset)
-40
-20
0
20
40
Time (min relative to start of furrow elongation)

**Figure S1. The start of chytrid cellularization coincides with the recruitment of nuclei to the plasma membrane and produces daughter cells of homogenous size, related to Figures 1 and 2. (A)** Distribution of sizes of chytrid zoospores (magenta) relative to artificial latex beads of 2 μm (teal). 95% of zoospores are within 0.57 μm of the mean (2.89 μm). $N_{zoospores}$ = 3785, $N_{beads}$= 7272. Measurements made with a Coulter counter. (**B)** Timing and duration from Stage 1 (magenta; vesicular furrow formation) to Stage 2 (green; furrow elongation) for each sporangium (row) relative to peak cortical nuclear migration. Different bar colors represent experiments from different days. (**C & D)** Cortical nuclear ratio (y-axis) relative to Stage 1 (C) or relative to Stage 2 (D). Note how the peak ratio tends to happen after Stage 1 and coincides with Stage 2. Same data as in A. Different bar colors represent experiments from different days.

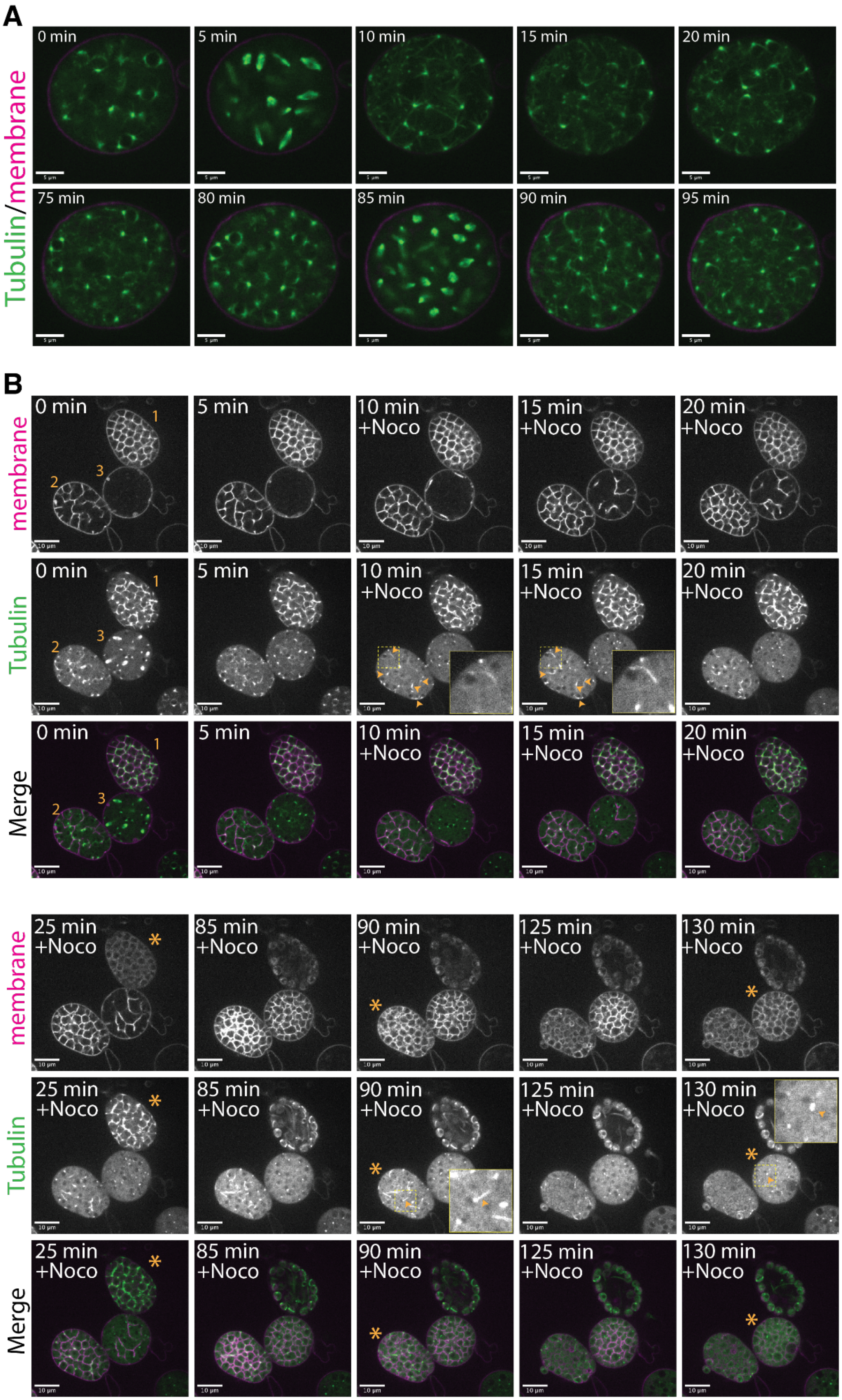
A
Tubulin/membrane
0 min
5 min
10 min
15 min
20 min
75 min
80 min
85 min
90 min
95 min
B
membrane
Tubulin
Merge
0 min
5 min
10 min +Noco
15 min +Noco
20 min +Noco
25 min +Noco
85 min +Noco
90 min +Noco
125 min +Noco
130 min +Noco
1
2
3
*

**Figure S2. Chytrid microtubule structures highlighted by the fluorescently-tagged alpha-tubulin and the effects of Nocodazole in these structures and ciliogenesis during cellularization, related to Figures 3 and 4. (A)** Nuclear-associated foci of fluorescently-tagged tubulin are likely chytrid centrosomes. Time-series Images of a sporangium during growth, before entering into cellularization. Expression of fluorescently-tagged chytrid alpha-tubulin highlights centrosome-like foci at the poles of each nucleus that turn into opposing spindle poles during nuclear division. Scale bar is 5 μm. **(B)** Nocodazole does not inhibit cellularization but produces defects in patterning and aciliated daughters. Example of three mother cells at different stages in cellularization during treatment with 2 μM Nocodazole (10 min onwards). Note how Nocodazole causes dissipation of fluorescent structures except for the bright foci at each nucleus (putative centrosome) and defects in polyhedral patterning. Mother 1 (see yellow numbers) is fully cellularized (Stage 4) with daughters fully ciliated. This mother completes cellularization (25 min; asterisk) and daughters escape the mother's cell wall. Mother 2 is at stage 2 of cellularization (branching & merging) and shows some nuclei starting to extend their axoneme (yellow arrows; 10 min) by the time of treatment. This results in a combination of daughters with and without cilia, but still completes cellularization (90 min; asterisk). Mother 3 is at stage 1-2 of cellularization (furrow initiation). Cellularization is completed at 130 min (asterisk) and produces daughters that completely lack cilia. Scale bar 10 μm.

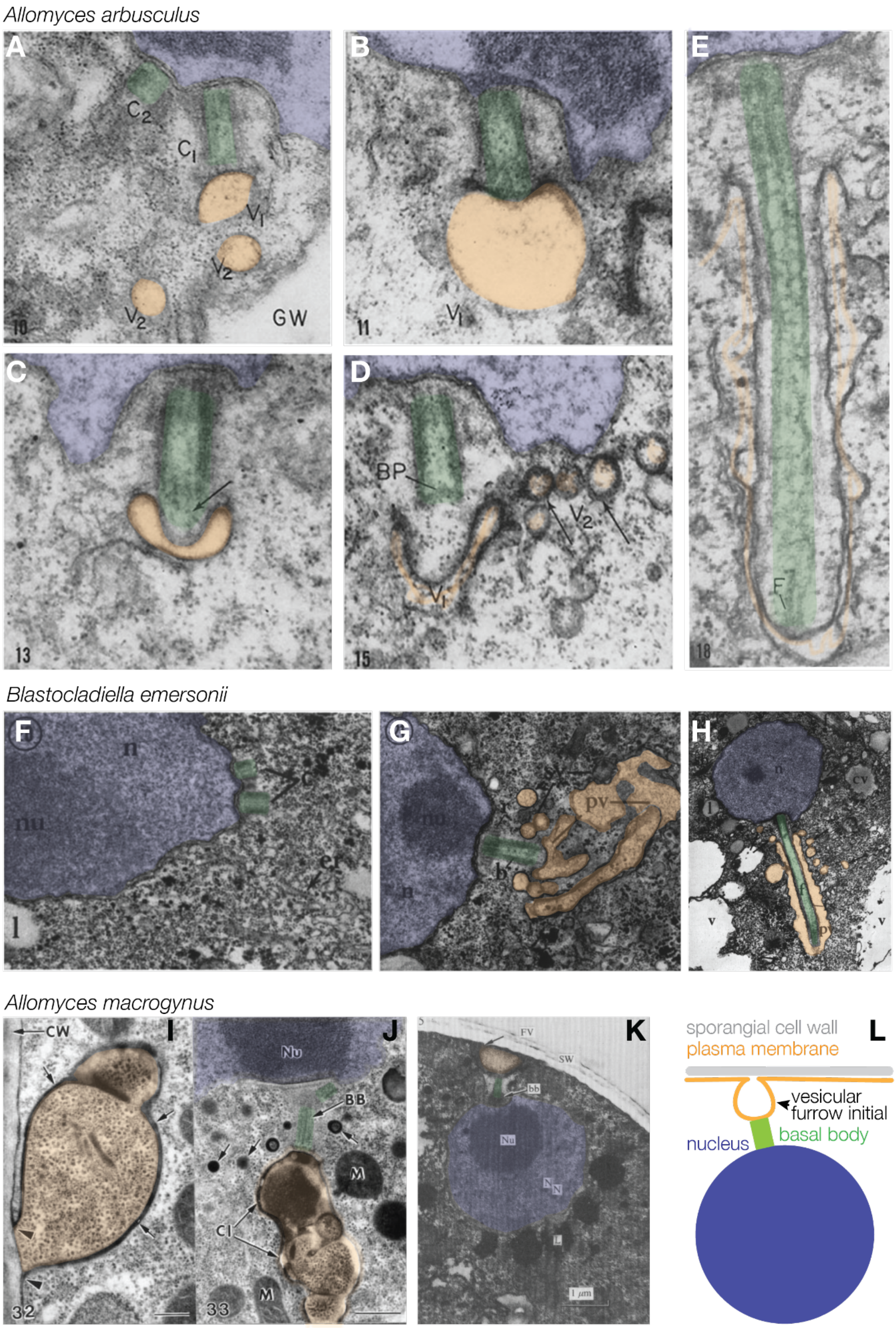
Allomyces arbusculus
A
C2
C1
V1
V2
V2
GW
10
B
V1
11
C
13
D
BP
V2
V1
15
E
F
18
Blastocladiella emersonii
F
n
nu
c
er
l
G
nu
n
pv
b
H
n
cv
v
v
Allomyces macrogynus
I
CW
32
J
Nu
BB
M
CI
M
33
K
FV
SW
bb
Nu
L
1 μm
L
sporangial cell wall
plasma membrane
vesicular
furrow initial
basal body
nucleus

**Figure S3. Ciliary vesicles form at nuclear-associated centrosomes during zoosporic fungi intracellular ciliogenesis and cellularization, related to Figure 3. A-E)** Intracellular ciliogenesis during cellularization in *Allomyces arbusculus:* **A)** recruitment of vesicles to the mother centriole. **B)** Formation of a ciliary vesicle at the distal end of the basal body. **C, D, & E)** The zoospore cilia are formed intracellularly; the axoneme extension is accompanied by the formation of a double membrane sheath. nucleus (pale blue), the centrioles, basal body, and axoneme (pale green), and associated membrane vesicles (pale yellow). Modified from images 10,11,13,15,18 from Renaud and Swift (1964).[S1] Original labels: C1: large centriole, C2: small centriole, V1: primary vesicle, V2: secondary vesicle, GW: gametangial wall, BP: basal plate, F: flagellar fibers. **F-H)** Intracellular ciliogenesis during cellularization in *Blastocladiella emersonii:* **F)** Nucleus before the start of cellularization. Note the centrioles tightly associated with the nuclear envelope. **G)** Recruitment of membrane vesicles to a ciliary vesicle at the distal end of the basal body. **H)** Intracellular axoneme and ciliary sheath extension. Modified from images 20-22 from Lessie and Lovett (1968).[S2] Original labels: C: centrioles, n: nucleus, nu: nucleolus, l: lipid droplet, er: rough reticulum, b: basal body, sv: secondary vesicle, pv: primary ciliary vesicle, cv: cleavage vesicle. **I-K)** Early cellularization in *Allomyces macrogynus:* **I)** membranous vesicular furrow initials form at the plasma membrane under the sporangial cell wall (CW). **J)** Cleavage element initials (CI) are intimately associated with the basal body (BB). K) The vesicular furrow initial, also referred to as the flagellar vesicle (FV) faces outwards from the distal end of the basal body (bb) and is continuous with the plasma membrane under the sporangium wall (SW). **I** and **J** are modified images 32 and 33 from Fisher, et al. (2000).[S3] Original Labels: CW: sporangia cell wall, Nu: nucleolus, BB: basal body, M: mitochondria, CI: Cleavage element initial. **K** is a modified image from Barron & Hill (1974).[S4] Original Labels: CW: sporangia cell wall, Nu: nucleolus, BB: basal body, M: mitochondria, CI: Cleavage element initial. FV: flagellar vesicle, SW: sporangial wall, Nu: nucleolus, N: Nucleus, L : Lipid. **L)** Our model for the formation of vesicular furrow initials in *Spizellomyces.* We propose that the "primary ciliary vesicles" in zoosporic fungi ciliogenesis literature are the same structures as our "vesicular furrow initials", playing a role in both ciliogenesis and cellularization by playing two roles simultaneously, tethering the nucleus to the future tubular furrow for cellularization and docking the basal body for axoneme extension during ciliogenesis.

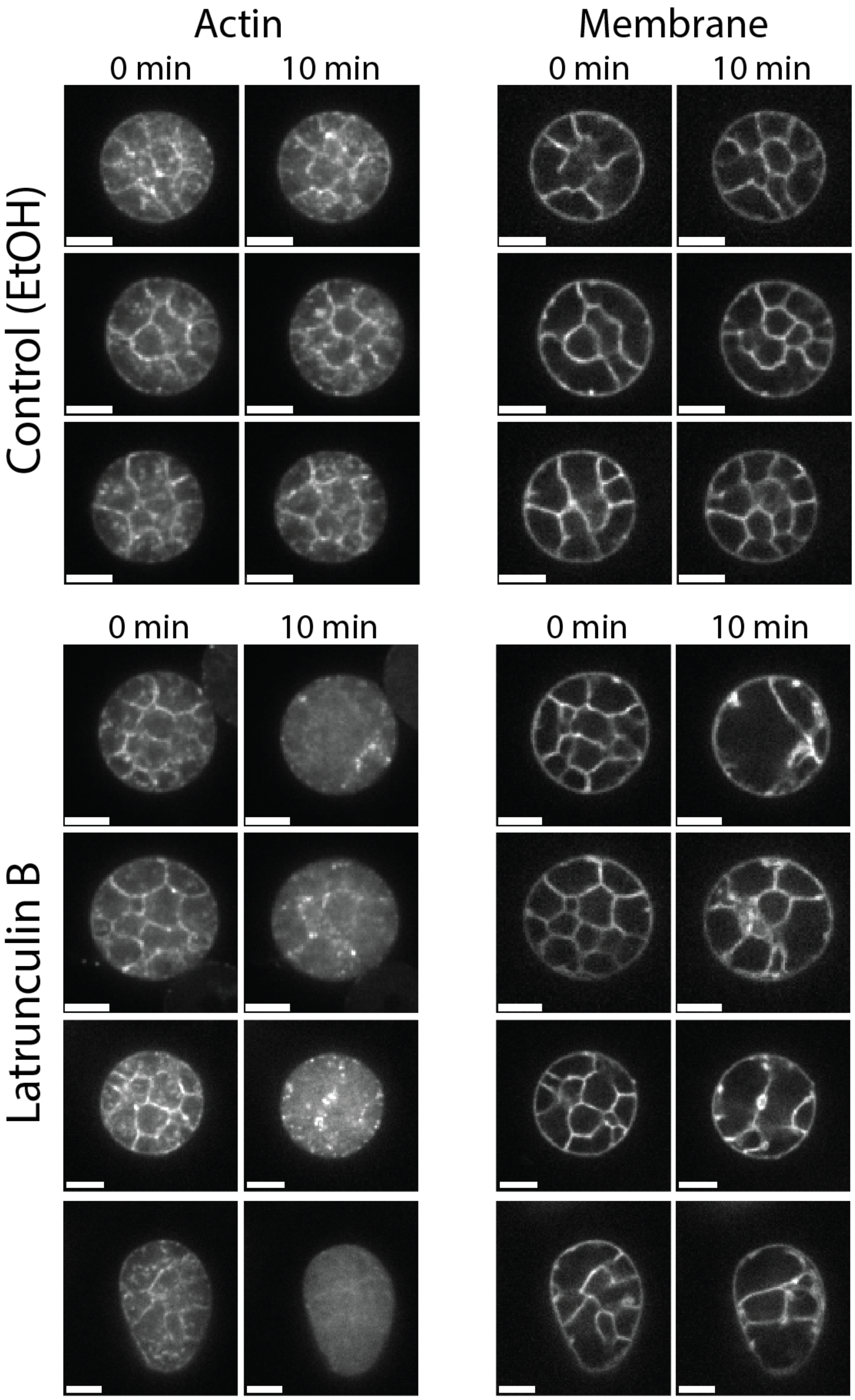


**Figure S4. Inhibition of actin polymerization with LatB causes arrest of cellularization and partial disassembly of the membrane cellularization network, related to Figure 5.** Panel shows a selection of sporangia in early cellularization from Figure 5A-B. The images shown correspond to the image acquired right before LatB is added (0 min), and 10 min of incubation in LatB, right before LatB washout. Note the slight increase in complexity of the membrane cellularization network in control mother cells compared to the disassembly of the cellularization networks in the cells treated with LatB. Scale bar 5 μm

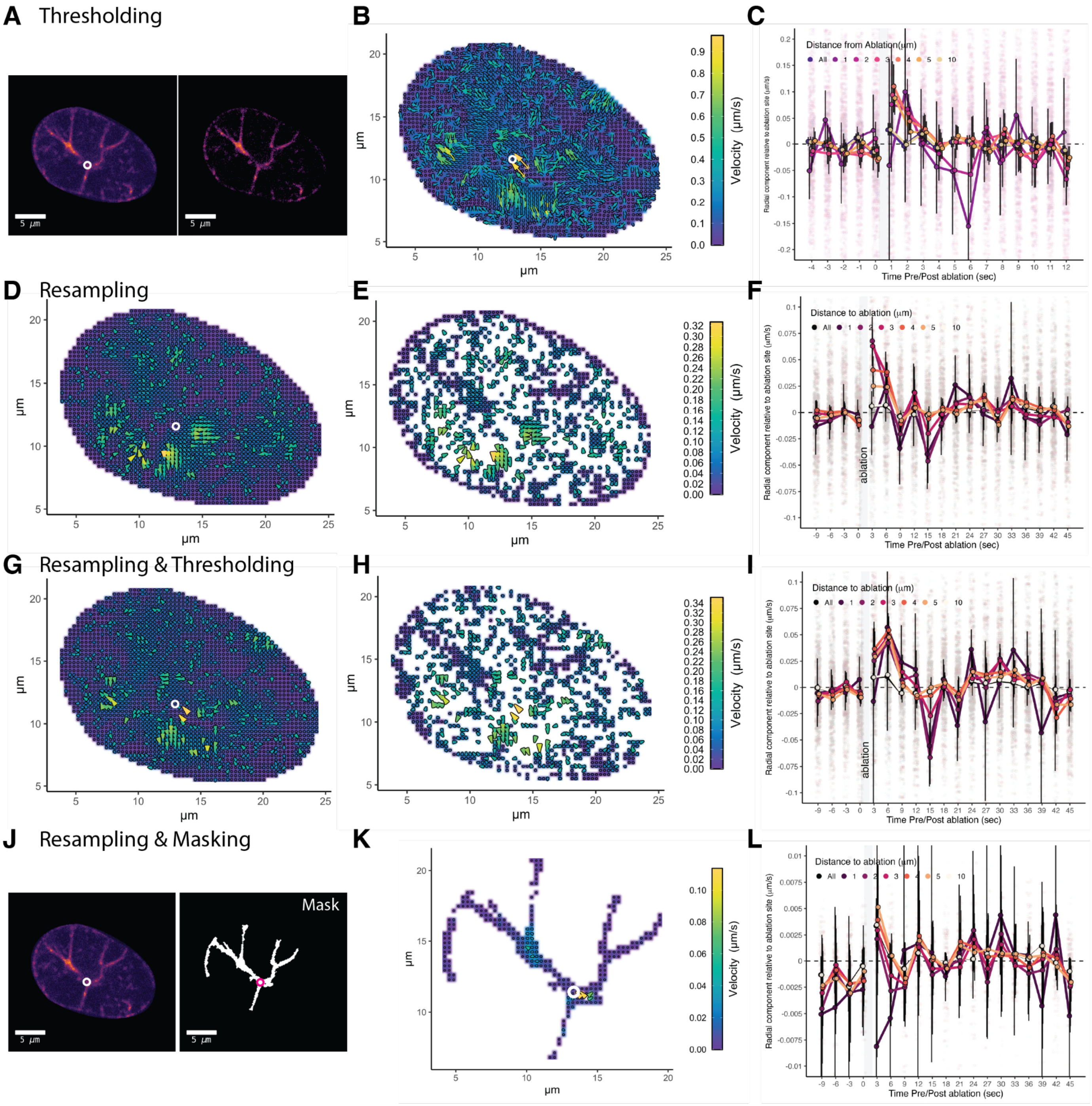


**Figure S5. The signal of recoil after ablation from PIV analysis is robust to different ways of filtering and thresholding the data, related to Figure 6. (A).** Example images before and after thresholding with Multi-Otsu. **(B)** Flow plot with local velocities estimated by PIV comparing the frame just before ablation (time 0) and the first frame after laser ablation (1 second from the end of ablation)—real time between images is 3 seconds, but time after the end of ablation is about 1 second. **(C)** Plot showing the radial component of velocities relative to the ablation site of all vectors with magnitude above zero and that pass the signal-to-noise test ('peak2peak'; threshold=0.5) from OpenPIV. Dots are mean values, and error bars show 99% confidence interval of the mean by bootstrap. For visualization purposes only, data within -0.2 and 0.2 μm/s is shown (42573 points; 85.6% of data). **(D)** Flow plot with local velocities estimated by PIV after resampling the image data every 3 seconds, such that the complete time series reflects the same sampling rate as before-after laser ablation. **(E).** Vectors from PIV that pass OpenPIV signal to noise test ('peak2peak'; threshold=0.5) and have a magnitude greater than zero are the ones used for estimating the radial component of velocities relative to the ablation site. **(F)** Plot showing the radial component of velocities relative to the ablation site for the resampled images. Dots are mean values, and error bars show 99% confidence interval of the mean by bootstrap. For visualization purposes only, data within -0.2 and 0.2 μm/s is shown (537754 points; 95% of data). **(G)** Flow plot with local velocities estimated by PIV after a combination of resampling and thresholding. **(H)** Vectors from (G) that pass signal to noise and have magnitudes greater than zero. **(I)** Plot showing the radial component of velocities relative to the ablation site for the resampled and thresholded images. Dots are mean values, and error bars show 99% confidence interval of the mean by bootstrap. For visualization purposes, only data within -0.2 and 0.2 μm/s is shown (49370 points; 92% of data). **(J)** Masking the resampled images to limit the PIV analyses to the furrow regions (based on thresholding of the sum of the images before ablation). **(K)** Flow plot with local velocities estimated by PIV comparing the frame just before ablation (time 0) and the first frame after laser ablation (3 seconds). **(L)** Plot showing the radial component of velocities relative to the ablation site for the resampled and masked images. Dots are mean values and error bars show 99% confidence interval of the mean by bootstrap. For visualization purposes only, data within -0.01 and 0.01 μm/s is shown (12042 points; 92% of data).

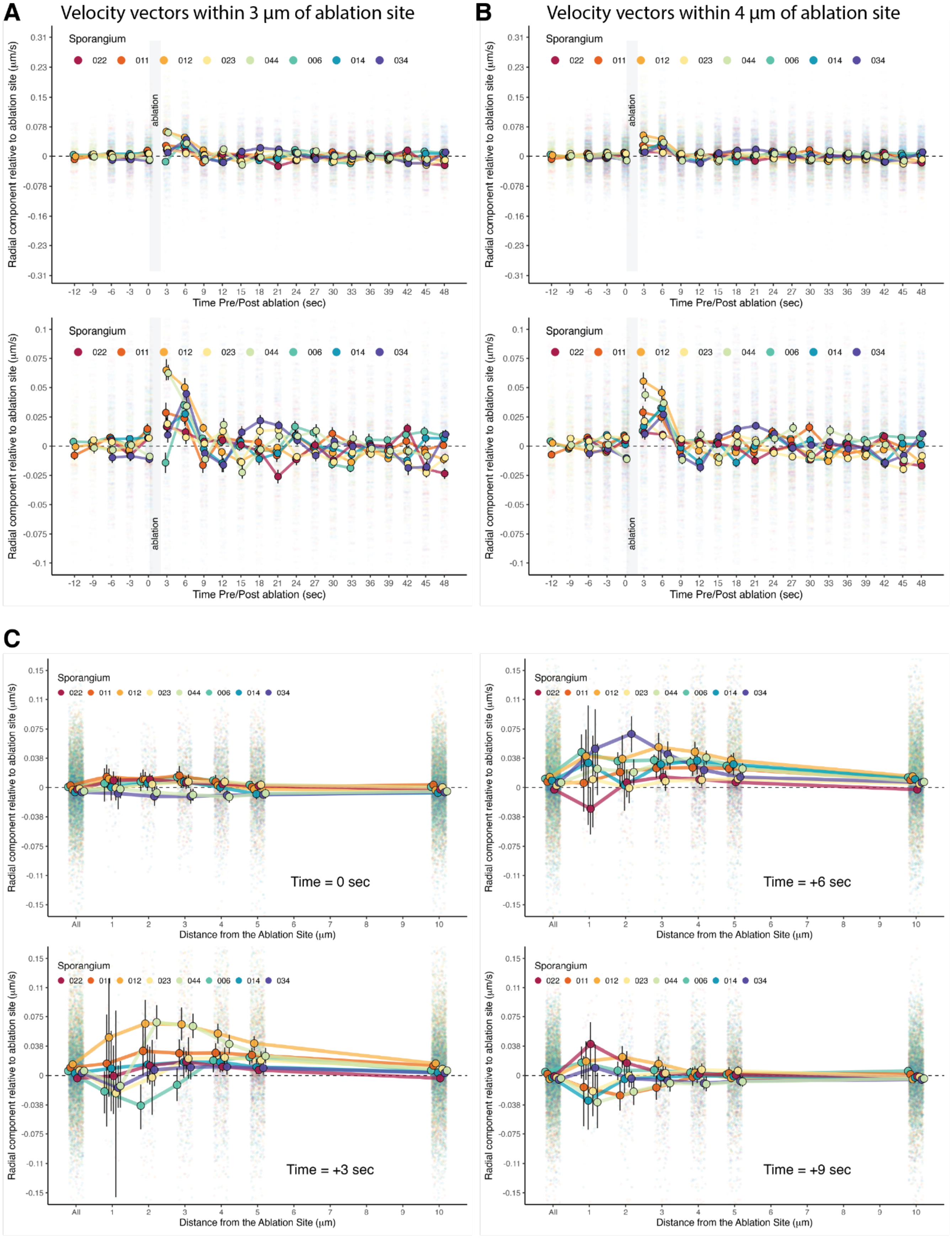


**Figure S6. Temporal dynamics, sampling, and distance effects from the ablation site on recoil velocities obtained via PIV, related to Figure 6. (A & B)** PIV retrieved high velocities within 3 µm and 4 µm from the ablation site when using data resampled every 3 seconds. **(A).** Plot showing the radial component relative to the ablation site (velocities away from—positive—or towards—negative—the ablation site) within 3 μm from the ablation site. The bottom shows an enlarged version of the top figure. For visualization purposes, we display data points within a range of -0.1 to 0.1 μm/s (this is 111009 points; 95% of the total data). **(B)** Plot showing the radial component relative to the ablation site (velocities away from—positive—or towards—negative—the ablation site) within 4 μm from the ablation site. This is the data shown in Figure 6 in the main text. Bottom panel shows enlarged version of top plot (Figure 6 data). Dots are mean values and error bars show 99% confidence interval of the mean by bootstrap. For visualization purposes, we display data points within a range of -0.1 to 0.1 μm/s (this is 176417 points; 95% of the total data). **(C)** Velocities away from or towards the ablation site at increasing distances from the ablation site during 3, 6, and 9 seconds after the beginning of ablation. Top. Plot showing the radial component relative to the ablation site (velocities away from—positive—or towards—negative—the ablation site) just before ablation. "All" uses all velocities within the sporangium that pass signal-to-noise OpenPIV test and have a magnitude greater than zero. The other plots show the distribution of velocities 3,6, and 9 seconds after ablation. Note that the greatest recoil velocities are detected between 2-4 μm from the ablation site and 3 seconds after ablation (4 seconds for sporangium 034 & 014). Dots are mean values and error bars show 99% confidence interval of the mean by bootstrap. For visualization purposes 25256 points (99%), 23198 points (97%), 24239 points (98%), and 24628 points (99%) are shown for each plot, respectively.

| Organism/Recombinant DNA | Strain/Name | Description | Accesion | Origin |
|---|---|---|---|---|
| *S. punctatus* | Lifeact-mClover3 | *Sp* transformed with T-DNA from pEM112C | EM112C | This study |
| | MLRC-mClover3 | *Sp* transformed with T-DNA from pEM110 | EM110 | This study |
| | mClover3-alpha-Tubulin | *Sp* transformed with T-DNA from pEM110 | EM102 | This study |
| | SAS6-mClover3 | *Sp* transformed with T-DNA from pEM120 | EM120 | This study |
| | RbNLS-mClover3 | *Sp* transformed with T-DNA from pGI3EM48 | EM48 | This study |
| | H2B-tdTomato | *Sp* transformed with T-DNA from pGI3EM20C | EM20C | Medina, et al. 2020 [S5] |
| *Agrobacterium* backbone pCAMBIA1300 | pEM112C | T-DNA insert: *H2Apr-Lifeact-mClover3-H2Ater:H2Bpr-P2A-T2A-hph-SynTer8** | pEM112C | This study |
| | pEM110 | *Sp*MRLC = SPPG_07268<br><br>T-DNA insert: *H2Apr-MRLC-mClover3-FLAG-MRLCter:H2Bpr-P2A-T2A-hph-SynTer8* | pEM110 | This study |
| | pEM102 | *AlphaTUB = Batrachochytium alpha-tubulin* (BDEG_00078)<br><br>T-DNA insert: *AlphaTUBpr-mClover3-BdenAlphaTUB-BdTUBter:H2Bpr-P2A-T2A-hph-SynTer8* | pEM102 | This study |
| | pEM120 | *Sp*SAS6 (SPPG_03341)<br><br>T-DNA insert: H2Apr-SAS6-mClover3-SAS6ter:H2Bpr-P2A-T2A-hph-SynTer8 | pEM120 | This study |
| *Backbone* pGI3EM22C[S5] | GI3EM48 | Lifeact-tdTomato replaced by RbNLS-mClover3; RbNLS = first 40aa of SPPG_07796<br><br>*H2Bpr-RbNLS-mClover3-ScADH1ter:H2Apr-hph-ScADH1ter* | pGI3EM48 | This study |

**SynTer8* is a short synthetic terminator (No 8) from Curran, et al. 2015[S6]

**Table S1. Strains and plasmids generated in this work. Related to STAR Methods**

**Supplemental References.**


S1. Renaud, F.L., and Swift, H. (1964). The Development of Basal Bodies and Flagella in *Allomyces arbusculus*. J. Cell Biol. *23*, 339–354. https://doi.org/10.1083/jcb.23.2.339.

S2. Lessie, P.E., and Lovett, J.S. (1968). Ultrastructural changes during sporangium formation and zoospore differentiation in *Blastocladiella Emersonii*. Am. J. Bot. *55*, 220–236.

S3. Fisher, K.E., Lowry, D.S., and Roberson, R.W. (2000). Cytoplasmic cleavage in living zoosporangia of *Allomyces macrogynus*. J. Microsc. *198*, 260–269. https://doi.org/10.1046/j.1365-2818.2000.00700.x.

S4. Barron, J.L., and Hill, E.P. (1974). Ultrastructure of Zoosporogenesis in *Allomyces macrogynus*. J. Gen. Microbiol. *80*, 319–327. https://doi.org/10.1099/00221287-80-2-319.

S5. Medina, E.M., Robinson, K.A., Bellingham-Johnstun, K., Ianiri, G., Laplante, C., Fritz-Laylin, L.K., and Buchler, N.E. (2020). Genetic transformation of *Spizellomyces punctatus*, a resource for studying chytrid biology and evolutionary cell biology. eLife *9*, e52741. https://doi.org/10.7554/eLife.52741.

S6. Curran, K.A., Morse, N.J., Markham, K.A., Wagman, A.M., Gupta, A., and Alper, H.S. (2015). Short Synthetic Terminators for Improved Heterologous Gene Expression in Yeast. ACS Synth. Biol. *4*, 824–832. https://doi.org/10.1021/sb5003357.
